# Molecular and functional dissection using CaMPARI-seq reveals the neuronal organization for dissociating optic flow-dependent behaviors

**DOI:** 10.1101/2025.06.04.654794

**Authors:** Koji Matsuda, Chung-Han Wang, Atsushi Toyoda, Tomoya Shiraki, Koichi Kawakami, Fumi Kubo

## Abstract

Optic flow processing is critical for the visual control of body and eye movements in many animals. Rotational and translational binocular optic flow patterns need to be clearly distinguished to induce different behavior outputs. However, the specific neuron types and their connectivity involved in this computation remain unclear. Here, we developed a method to link the functional labeling using a photoconvertible calcium indicator called CaMPARI2 and single-cell RNA-sequencing (CaMPARI-seq) to investigate the transcriptional profile of the pretectum, a center for processing optic flow in larval zebrafish. Using this technique, we identified a pretectal cluster expressing *tcf7l2*, which can be further classified into molecularly distinct subclusters. In vivo calcium imaging and cell ablation revealed that *nkx1.2lb*-positive pretectal neurons are commissural inhibitory neurons required for the optomotor response but not for the optokinetic response. Our genetic and functional dissection using CaMPARI-seq uncovered the neuronal organization essential for dissociating different optic flow-dependent behaviors.

## Main

Optic flow, the visual motion resulting from animals’ movements, plays an important role in estimating self-motion in invertebrates and vertebrates (Masseck and Hoffmann, 2009; Srinivasan and Zhang, 2000). Optic flow triggers the stereotyped compensatory responses of the eye (the optokinetic response; OKR) and body (the optomotor response; OMR) by which animals adjust and minimize self-motion-induced displacement (Brockerhoff et al., 1995; Muto et al., 2005; Neuhauss et al., 1999). Previous studies conducted in mammals and teleosts have demonstrated that the activation of the pretectum (a part of the accessory optic system in mammals) is sufficient to evoke OKR, whereas lesions or experimental inactivation suppress this behavior (Cazin et al., 1980; Kubo et al., 2014; Schiff et al., 1988). The afferent and efferent connections of the pretectum in mammals and teleosts suggest that the pretectum receives direct inputs from the retinal ganglion cells (RGCs) in the contralateral retina and transmits information to the brainstem and other structures in support of visual-oculomotor control (Giolli et al., 2006; Vanegas and Ito, 1983; Yáñez et al., 2018). In the zebrafish, a subset of RGCs that encode the direction of motion, namely the direction-selective (DS-) RGCs, project to a neuropil region of the pretectum (Baier and Wullimann, 2021; Kramer et al., 2019; Yildizoglu et al., 2020). This retinorecipient neuropil, primarily in the arborization field AF5, is innervated by optic flow-responsive pretectal neurons (Kramer et al., 2019). A subset of the pretectal neurons projects to downstream premotor/motor areas such as the nucleus of the medial longitudinal fasciculus (nMLF) in the midbrain and reticular formation in the hindbrain, which produce motor output (Kramer et al., 2019; Orger et al., 2008).

Previous systematic and unbiased functional imaging studies in zebrafish have demonstrated that hundreds of DS neurons in the pretectum respond to optic flow (Kubo et al., 2014; Naumann et al., 2016; Portugues et al., 2014). Furthermore, these DS pretectal neurons show diverse physiological properties (reviewed in Matsuda and Kubo, 2021), including preferred directions of motion (Wang et al., 2019), receptive field size and location (Wang et al., 2020), motion sensitivity defined by higher-order correlations (Yildizoglu et al., 2020), binocular integration (Kubo et al., 2014; Naumann et al., 2016; Wang et al., 2019; Zhang et al., 2022), and temporal integration (Dragomir et al., 2020). Among them, binocular integration of optic flow is of high behavioral relevance. In lateral-eyed animals, including the zebrafish, comparison of the motion information between the left and right eyes is an integral step in estimating optic flow patterns across a wide area of the visual field (Krapp et al., 2001; Kubo et al., 2014; Naumann et al., 2016; Portugues et al., 2014; Wylie et al., 1998). Functional imaging has demonstrated that a large population of zebrafish pretectal neurons integrate binocular optic flow (Kubo et al., 2014; Naumann et al., 2016; Portugues et al., 2014). Notably, most of these binocular neurons selectively respond to translational optic flow, and rotation-selective neurons comprise only a minor part (Kubo et al., 2014; Zhang et al., 2022). These observations led to the idea that binocular computation in the pretectum is suited for processing translation-induced optic flow but not rotation-induced optic flow, and this property ensures the appropriate directionality of OMR and OKR behaviors to be unambiguously encoded at the level of the pretectum (Zhang et al., 2022). However, the specific neuron types and connectivity enabling this separate processing remain unclear.

Despite extensive characterization of the physiological properties of the optic flow processing circuit, the causal link between the identified neuron types and behavior remains largely unknown. This is partly due to the unavailability of techniques to manipulate optic flow-responsive neurons in a subtype-specific manner. To overcome this problem, we sought to genetically label and manipulate different functional subtypes of pretectal neurons. Vertebrate pretectum is derived developmentally from an evolutionarily conserved segment of the diencephalon known as prosomere 1 (p1) (Brożko et al., 2022; Puelles and Rubenstein, 2003; Virolainen et al., 2012; Wullimann et al., 2024). Although some transcription factors that are expressed by the p1 domain have been described, current knowledge about cell-type-specific molecular signatures within the pretectum is limited. A recent study conducted using high-throughput single-cell RNA sequencing (scRNA-seq) revealed the molecular taxonomies of zebrafish visual neurons, including pretectal neurons (Sherman et al., 2023). However, it is challenging to associate individual transcriptomic types with their function, behaviors, and morphological connectivity.

In this study, we employed transcriptional profiling of pretectal neurons linked with visual functions. We strategically combined activity-dependent fluorescent labeling by CaMPARI and the scRNA-seq technique to molecularly classify optic flow-responsive neurons. We identified a transcriptomic cell type that contained optic flow-responsive cells in the pretectum, which we further classified into seven distinct transcriptomic cell types. Using CRISPR/Cas9 genome editing and intersectional strategies, we established a set of transgenic lines to genetically access several molecularly defined pretectal cell types. We identified two molecularly distinct subtypes of pretectal neurons that were morphologically distinct and composed of different sets of functional response types. We found that a subset of pretectal neurons that express the transcription factor *nkx1.2lb* are inhibitory commissural neurons that are required for the optomotor response but dispensable for the OKR. Overall, our functionally targeted transcriptional profiling identified the circuit organization and mechanisms required for computing behaviorally relevant information during optic flow processing.

## Results

### Molecular profiling of functionally identified pretectal neurons using CaMPARI-seq

To obtain genetic access to the functionally diverse types of optic-flow responsive pretectal neurons, we employed single-cell RNA sequencing of functionally identified neurons, which we termed CaMPARI-seq (Fig. 1a). To specifically isolate optic flow-responsive cells, we fluorescently labeled them using CaMPARI2, a Ca^2+^-dependent photoconvertible protein that changes its fluorescence from green to red by UV illumination, thereby labeling neurons that are activated only during a specific temporal window (Extended Data Fig. 1a) (Fosque et al., 2015; Moeyaert et al., 2018). We generated a transgenic zebrafish line *Tg(elavl3:NLS-CaMPARI2)* expressing CaMPARI2 fused with the nuclear localization signal (NLS-CaMPARI2) in most neurons. We reasoned that the conventional cytosolic CaMPARI2 would not be ideal because the cytosolic CaMPARI2 signal distributed in the neurites would be eliminated during the following cell dissociation procedure required for the scRNA-seq, attenuating the CaMPARI2 signals in the dissociated cells. To circumvent this, we fused CaMPARI2 with the nuclear localization signal (NLS) so that the NLS-CaMPARI2 protein localized in the soma would not be lost during cell dissociation. Similar to the previously reported cytosolic CaMPARI in the zebrafish brain (Fosque et al., 2015), UV illumination at 405 nm induced the robust photoconversion of NLS-CaMPARI2 in response to the proconvulsive compound 4-aminopyridine (4-AP) (Extended Data Fig. 1b).

**Fig. 1:**
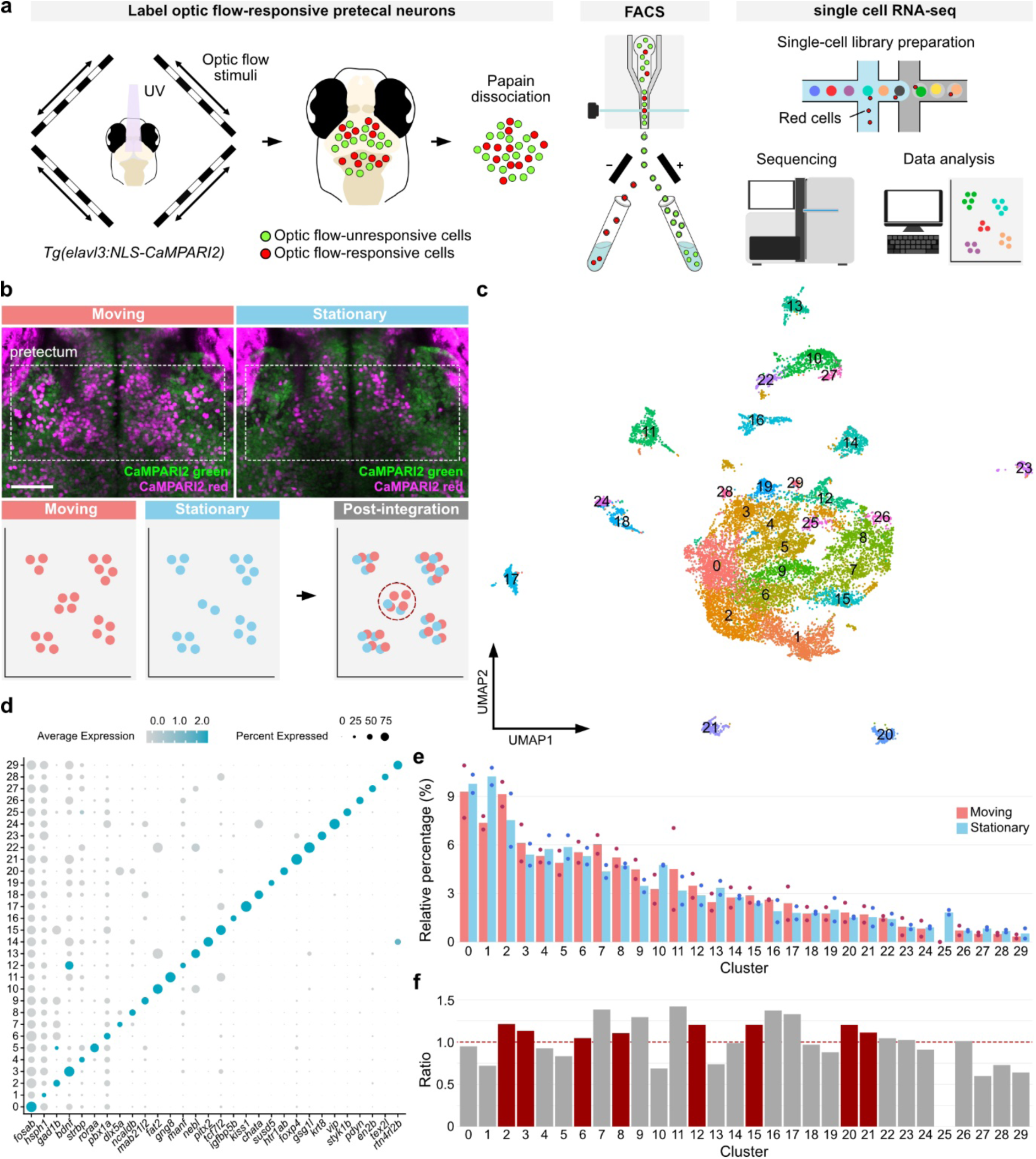
Molecular profiling of functionally labeled pretectal neurons using CaMPARI-seq. **a,** Experimental workflow for identifying marker genes for optic flow-responsive pretectal neurons. **b,** (Top) Moving and stationary conditions used for CaMPARI-seq. Note that a greater number of CaMPARI2 red cells are detected in the “moving” condition compared to the “stationary” condition. Scale bar, 50 μm. (Bottom) Ratio of cells derived from “moving” datasets over those from “stationary” datasets (MS index). **c,** UMAP embedding of all sequenced cells after integrating two “moving” datasets and two “stationary” datasets. **d,** Markers for clusters. Color shade represents the average expression level within a cluster (average expression). Dot size represents the percentage of cells expressing the marker in a cluster (percent expressed). **e,** Relative frequency (y axis) of each cluster (x axis), ordered from highest to lowest. Each data point represents one dataset and their average is shown as bar graph. **f,** Ratio of cell frequency of “moving” datasets over that of “stationary” datasets for each cluster (MS index). Red clusters with MS index higher than 1 were considered as candidates for the pretectal optic flow-responsive clusters (see text for details).

To photoconvert NLS-CaMPARI2 during optic flow stimulation, we illuminated *Tg(elavl3:NLS-CaMPARI2)* larvae with UV LED coupled with the presentation of a visual stimulus that consisted of four binocular optic flow patterns (horizontally moving gratings); namely, clockwise, counter-clockwise, forward, and backward motions, referred to as the “moving” experiment (Fig. 1a and Extended Data Fig. 1a). As a control, we used stationary gratings, referred to as the “stationary” experiment. During “moving” visual stimulation, neurons in the pretectum and its surrounding brain areas (such as tectum, thalamus, habenula, cerebellum, etc.) of *Tg(elavl3:NLS-CaMPARI2)* larvae were photoconverted by UV light (Fig. 1b and Extended Data Fig. 1c). Consistent with the previously reported spatial distribution of optic flow-responsive cells (Kubo et al., 2014; Naumann et al., 2016; Portugues et al., 2014; Wang et al., 2019), photoconverted CaMPARI2-red neurons were enriched in the middle part of the pretectum, as well as the lateral pretectum; i.e., the pretectal migrated area M1, in the moving experiments. Significantly fewer neurons were photoconverted in these regions in the stationary experiments, indicating that the neurons that responded to the optic flow were reliably labeled (Fig. 1b and Extended Data Fig. 1c). After the photoconversion during optic flow stimulation, we collected CaMPARI2-red neurons by FACS and transcriptionally profiled them using single-cell RNA-seq. Our transcriptional profiling was not designed to capture activity-dependent transcriptional changes in response to visual stimulation, since the transcriptomic analyses were performed after the visual stimulation and photoconversion (typically minutes to hours later), by which time transiently expressed immediate early genes are already turned off. We obtained two replicate datasets from the “moving” condition and two others from the “stationary” control and integrated all datasets into a single dataset by the SCTransform approach (Extended Data Fig. 1d-g and Methods). Clustering of the integrated dataset yielded 30 clusters based on the expression of the marker genes (Fig. 1c,d). To identify transcriptomic clusters in which optic flow-responsive cells were enriched, we focused on cell clusters that contained more abundant cells detected in the datasets obtained from the “moving” condition compared to those from the “stationary” condition (Fig. 1b). For each cluster, we calculated the fold change in the proportion of cells between the “moving” and “stationary” conditions (the MS index; see the Methods section for details). If the proportion of cells in the “moving” condition is higher than that in the “stationary” condition, the MS index would be greater than 1. Among the 16 clusters that showed an MS index higher than 1, we excluded eight since their identities were unambiguously annotated based on published data (Extended Data Fig. 1h, see the Methods section for details). The remaining eight clusters were considered candidate clusters that potentially contained optic flow-responsive cells in the pretectum (Fig. 1e,f).

### CaMPARI-seq identifies pretectal neuron subtypes labeled by a set of genes including *tcf7l2* and *gad1b*

Subsequently, we performed reclustering of the cells using the eight pretectum candidate clusters, resulting in 16 clusters that were identifiable by one or a few specific marker genes (Fig. 2a,b). After this round of clustering, two clusters showed a high MS index, suggesting enrichment of the optic flow-responsive cells derived from the “moving” dataset compared with the “stationary” dataset (Fig. 2c,d). Close inspection of one of the two candidate clusters (cluster 7) suggested that it corresponds to a putative hypothalamus region, based on the expression of *sim1a* and *fezf2* (Wolf and Ryu, 2013). The other cluster (cluster 5) was characterized by the expression of *tcf7l2 (transcription factor 7-like 2)*, which has been shown to be expressed in the broad areas of the diencephalon (e.g., the thalamus, habenula, etc.) including the pretectum in early zebrafish larvae (2 dpf) (Brożko et al., 2022). Next, we sought to clarify if this cluster corresponds to the pretectum but not the other regions of the diencephalon. We used a combination of genes whose expression domains delineate boundaries for the pretectum, thalamus, and prethalamic areas in the larval zebrafish brain (Wullimann et al., 2024). A marker gene for the pretectum, *pax7a,* was expressed in cluster 5, whereas the thalamus marker gene, *lhx9*, was not detectable in cluster 5 and was weakly expressed in cluster 1 (Fig. 2e,f). Furthermore, we examined known marker genes for the P1 domain in the mouse embryonic diencephalon and prenatal pretectum, such as *tal2* and *gata3* (Virolainen et al., 2012), and found that both *tal2* and *gata3* were expressed in cluster 5 (Fig. 2b,e,f). These expression patterns of a combination of genes suggest that cluster 5 corresponds to the pretectum. A similar set of genes (i.e., *tcf7l2*, *pax7a*, *tal2*, and *gata3*) were also expressed in cluster 15, which was marked by the expression of an additional gene, tyrosine *hydroxylase* (*th)*, likely corresponding to dopaminergic neurons in the pretectum (Fig. 2b,f). We excluded this cluster from further analyses because of its low MS index (Fig. 2c,d). To determine whether *tcf7l2* and *tal2* are expressed in the pretectum at the late larval stage (5–6 dpf), we generated gene-specific Gal4 driver lines using CRISPR/Cas9-mediated genome integration and crossed them with the *Tg(UAS:GFP)* reporter line (Fig. 2g–j). Similar to the expression pattern of *tcf7l2* in early zebrafish embryos (Brożko et al., 2022), *Tg(tcf7l2-hs:Gal4FF)* labeled cells in the pretectum and other brain regions of diencephalic origin, as well as parts of the tectum and forebrain (Fig. 2g). In the pretectum, *Tg(tcf7l2-hs:Gal4FF)*-labeled cells are tightly packed, indicating that the majority of cells expressed *tcf7l2* (Fig. 2h). Immunostaining of Tcf7l2 confirmed that the majority of the cells in the pretectum expressed Tcf7l2 (Fig. 2k–n). *Tg(tal2-hs:Gal4FF);Tg(UAS:GFP)* labeled cells in the pretectum and tectum (Fig. 2i,j). Compared with the ubiquitous expression of *tcf7l2*, *Tg(tal2-hs:Gal4FF)*-labeled cells were more sparsely distributed in the pretectum. To determine whether cluster 5 neurons are excitatory or inhibitory, we examined the expression of *slc17a6b* (AKA. *vglut2a,* the marker for glutamatergic excitatory neurons) and *gad1b* (the marker for GABAergic inhibitory neurons). Interestingly, the majority of cells (80.47%) in this cluster expressed *gad1b* and only 5.73% of them expressed the glutamatergic marker gene *slc17a6b* (Fig. 2e,f). Taken together, our CaMPARI-seq technique identified the transcriptional profile of putative optic flow-responsive GABAergic cells in the pretectum and obtained genetic access to them.

**Fig. 2:**
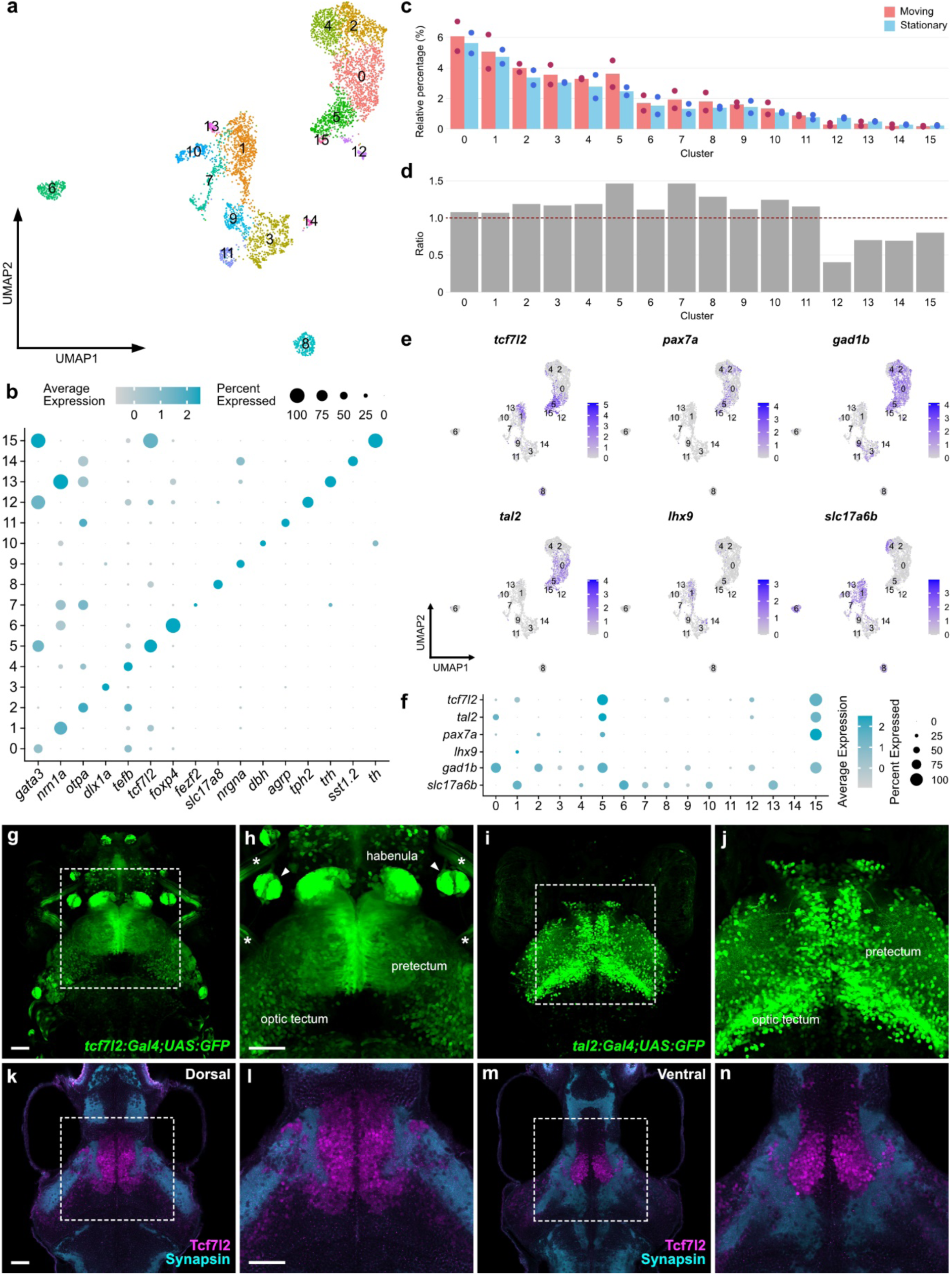
Identification of optic flow-responsive pretectal cluster and marker genes. **a,** UMAP embedding of pretectal candidate cells. Cells from two “moving” datasets and two “stationary” datasets are integrated and plotted. **b,** Markers for clusters. Color shade represents the average expression level within a cluster (average expression). Dot size represents the percentage of cells expressing the marker in a cluster (percent expressed). **c,** Relative frequency (y axis) of each cluster (x axis), ordered from highest to lowest. Each data point represents one dataset and average is shown as bar graph. **d,** Ratio of cells derived from “moving” datasets over those from “stationary” datasets (MS index). **e,** Gene expression plots of cells embedded in UMAP space. **f,** Expression of *tcf7l2*, *tal2*, *pax7a*, *lhx9*, *gad1b*, *slc17a6b* for each cluster. Color shade represents the average expression level within a cluster (average expression). **g-j,** Maximum z-projection of a substack from *Tg(tcf7l2-hs:Gal4FF);Tg(UAS:GFP)* larva (**g**,**h**) and *Tg(tal2-hs:Gal4FF);Tg(UAS:GFP)* larva (**i**,**j**). Asterisks depict extraocular muscles. Arrowheads indicate lateral line neuromasts. **k-n,** Immunostaining of Tcf7l2 co-stained with Synapsin in the dorsal (**k**,**l**) and ventral (**m**,**n**) pretectum. Scale bars, 50 μm.

### *tcf7l2*+ neurons label most of the optic flow-responsive cells in the pretectum

Next, we investigated whether the *tcf7l2*-positive pretectum cluster corresponds to the optic flow-responsive neurons. We tested if *tcf7l2*+ neurons responded to optic flow stimuli by two-photon Ca^2+^ imaging using *Tg(tcf7l2-hs:Gal4FF);Tg(UAS:GCaMP6s)* larvae at 6 dpf. Visual stimulation was presented from a 360-degree arena, consisting of eight different combinations of monocular and binocular horizontal gratings (Kubo et al., 2014, Fig. 3). We found that *tcf7l2*-positive neurons, particularly those located near the posterior boundary of the *tcf7l2-*expressing domain, strongly responded to the optic flow stimuli (Fig. 3a,d). When stimulated with a series of monocular and binocular motions, pretectal neurons in the zebrafish larvae can be classified into different functional types based on their binary (On or Off) response to each of the eight different motion phases (Kubo et al., 2014). Of the 256 (2^8^) theoretically conceivable binary response types, monocular direction-selective and translation-selective response types are frequently represented in the pretectum using the pan-neuronal GCaMP line *Tg(elavl3:H2B-GCaMP6s)* as previously described (Kubo et al., 2014, Extended Data Fig. 2). The application of the same classification to the response profiles of *tcf7l2*-positive neurons revealed that most of these major response types were highly represented (Fig. 3a–c). Furthermore, the spatial distribution of optic flow-responsive cells in the pretectum of the *Tg(tcf7l2-hs:Gal4FF);Tg(UAS:GCaMP6s)* larvae revealed that monocular DS cells are located contralateral to the visually stimulated eye in most cases (e.g., neurons responding to motion in the right eye are located in the left brain), whereas translation-selective neurons (especially the more abundant BEL and BER types) are located in both pretectal hemispheres (Fig. 3d). A similar spatial organization was observed for *Tg(elavl3:H2B-GCaMP6s)* larvae (Extended Data Fig. 2). These results suggest that *tcf7l2* labeled most optic flow-responsive neurons in the pretectum. This observation is consistent with the uniform distribution of *tcf7l2* expression within the pretectum (Fig. 2g,h,k–n).

**Fig. 3:**
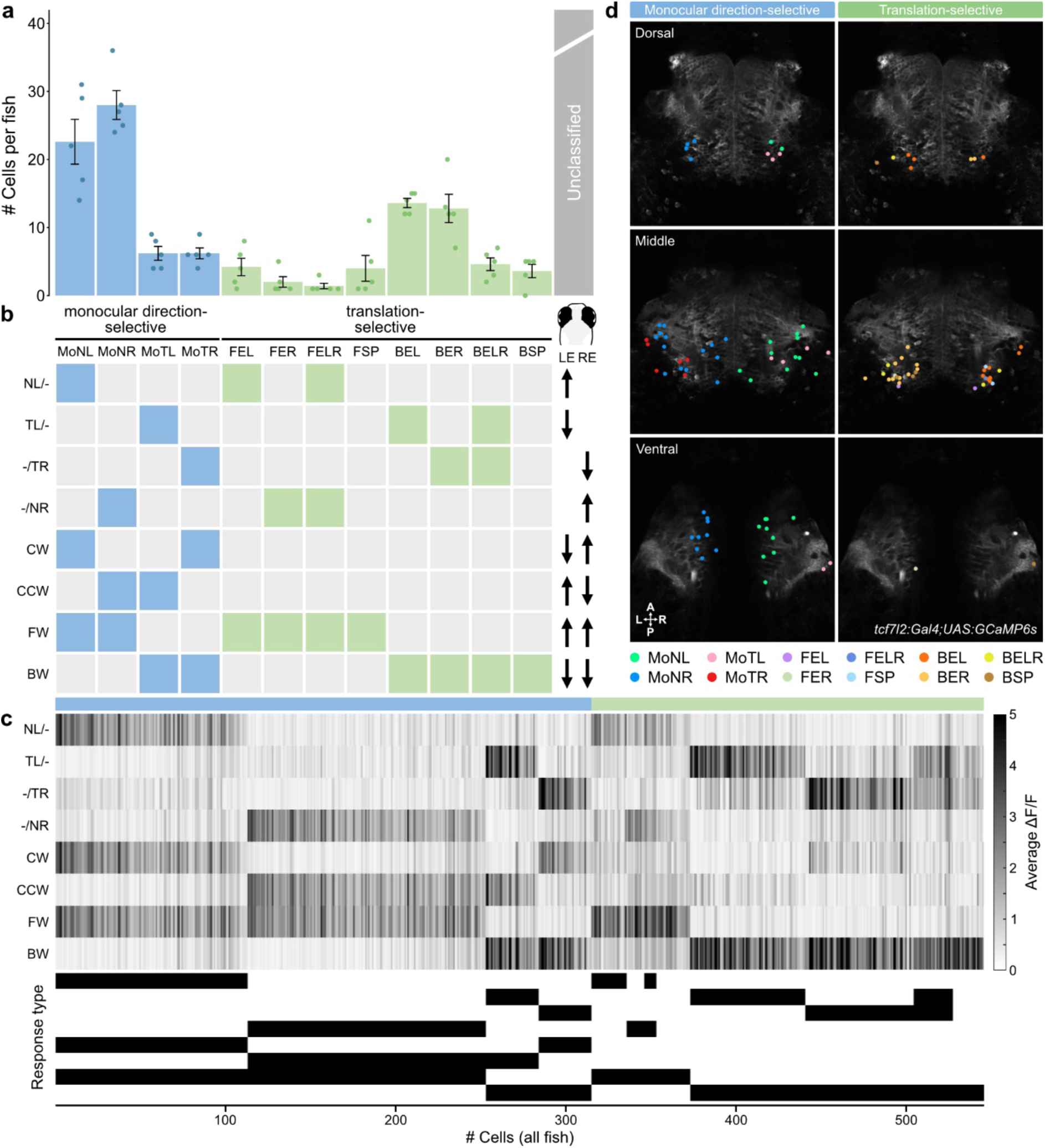
*tcf7l2*+ neurons in the pretectum respond to optic flow. **a,** A histogram of the frequent response types (“simple”, blue, “translation-selective”, green) detected in *Tg(tcf7l2-hs:Gal4FF);Tg(UAS:GCaMP6s)* larvae. The remaining response types were listed as unclassified (gray). Each bar represents the average # of cells per fish. Error bars represent ±SEM (*n* = 5 fish). **b,** Nomenclature of the visual stimulus protocol and response types as previously described (Kubo et al., 2014). **c,** (Top) Raster plot showing the responses to the eight stimulus phases (*n* = 546 cells, pooled from 5 fish). Cells are ordered according to their correlation coefficient to the corresponding regressor (within each response type). ΔF/F values across the 3 repetitions of the visual stimulation were averaged. (Bottom) Binary response type barcode of each cell. **d,** Spatial distribution of the response types (as shown in (**a**)) in a representative *Tg(tcf7l2-hs:Gal4FF);Tg(UAS:GCaMP6s)* fish. Cells are color-coded according to their response type. Average image of GCaMP6s signal of the dorsal, medial and ventral representative planes of the pretectum is used as a background. For the middle and ventral planes, cells detected in the same plane, as well as those detected in 10 µm above and 10 µm below the corresponding plane, are plotted. For the dorsal plane, cells detected in the same plane, as well as those detected in 10 µm above the corresponding plane, are plotted. For the data from all individual planes, see Extended Data Fig. 4 and 5. A, anterior; P, posterior; L, left; R, right.

### Pretectal optic flow neurons can be further clustered into subsets

Since *tcf7l2* marks the majority of optic flow-responsive neurons in the pretectum, we asked if the functional subtypes of the pretectum could be identified by further clustering of the *tcf7l2*-positive pretectal cluster (i.e., cluster 5 in Fig. 2). We reclustered cells contained in the *tcf7l2*-positive pretectal cluster, which yielded seven subclusters (Fig. 4a,b,e). The MS index of each cluster revealed that all clusters had a higher contribution from the “moving” dataset compared to the “stationary” one to varying degrees (Fig. 4c,d). We examined the expression pattern of the marker genes of different subclusters by hybridization chain reaction (HCR) RNA-FISH. Most genes were sparsely expressed in the pretectum area delineated by the expression of *tcf7l2* (Fig. 4f and Extended Data Fig. 3). For example, *nkx1.2lb* (NK1 transcription factor related 2-like, b) (Bae et al., 2004) was expressed in the dorsal-posterior part of the pretectum, while *mafaa* (MAF bZIP transcription factor Aa) was expressed in the ventral-lateral part of the pretectum (Fig. 4g‒q). *npy* (*neuropeptide Y*) was expressed more anteriorly compared with *nkx1.2lb*, while *penkb* was expressed more medially compared with *mafaa* (Extended Data Fig. 3). Notably, in most cases, the expression patterns of different marker genes identified from different clusters did not spatially overlap, suggesting that the transcriptional subtypes correspond to anatomically distinct cell types. As expected, for all of the clusters, the cells were largely *gad1b*+ GABAergic, while only a minority of the cells were *slc17a6b*+ glutamatergic (Fig. 4b).

**Fig. 4:**
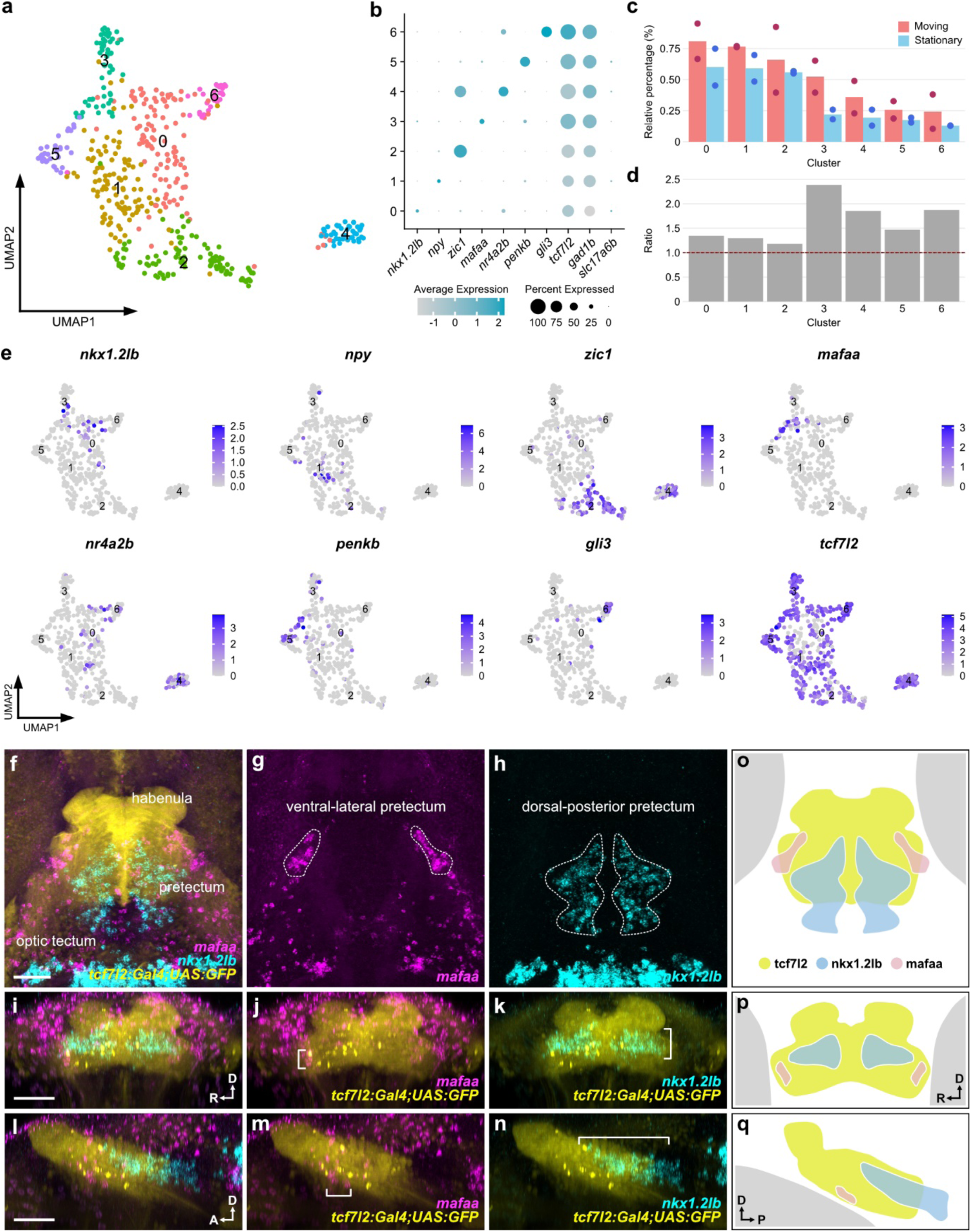
Pretectal neurons consists of diverse molecular types. **a,** UMAP embedding of pretectal subtypes. Cells from two “moving” datasets and two “stationary” datasets are integrated and plotted. **b,** Markers for clusters. Color shade represents the average level of marker expression in a cluster (average expression). Dot size represents the percentage of cells expressing the marker in a cluster (percent expressed). **c,** Relative frequency (y axis) of each cluster (x axis), ordered from highest to lowest. Each data point represents one dataset and average is shown as bar graph. **d,** Ratio of cells derived from “moving” datasets over those from “stationary” datasets (MS index). **e,** Gene expression plots of cells embedded in UMAP space. **f-n,** Substack maximum z-projections of double HCR-FISH stains of *mafaa* and *nkx1.2lb* in *Tg(tcf7l2-hs:Gal4FF);Tg(UAS:GFP)* larva in the dorsal (**f**-**h**), coronal (**i**-**k**), and sagittal (**l**-**n**) views. **o-q,** Schematic representation of the expression patterns in the pretectum in the dorsal (**o**), coronal (**p**) and sagittal (**q**) views. D, dorsal; A, anterior; R, right; P, posterior. Scale bars, 50 μm.

Genetic targeting of *mafaa-* and *nkx1.2lb-*positive neurons identified anatomically and functionally distinct pretectal subtypes

To determine whether transcriptional subtypes correspond to different anatomical features and/or functional subtypes of the optic flow-responsive pretectal neurons, we focused on the following two subclusters: *mafaa*+ and *nkx1.2lb*+ clusters. First, to label *mafaa*-positive cells, we generated a Gal4 knock-in transgenic line of *mafaa* gene *Tg(mafaa-hs:Gal4FF)*. As expected from the HCR RNA-FISH data (Fig. 4f,g), neurons labeled in *Tg(mafaa-hs:Gal4FF);Tg(UAS:GCaMP6s)* were located in the lateral part of the pretectum, the pretectal migrated area M1, partially homologous to the mammalian accessory optic system (Sherman et al., 2023; Wu et al., 2020) (Fig. 5a,b). The projection of these *mafaa*-positive neurons remained close to the soma and did not extend outside of this local region, suggesting that they extended local projections within a neuropil region near their cell bodies (Fig. 5c‒e). Furthermore, the neurites of *mafaa*+ neurons overlapped with the AF5 and AF6 of the retinal ganglion cell (RGC) axon terminals, where direction-selective RGCs innervate (Kramer et al., 2019; Yildizoglu et al., 2020) (Fig. 5f‒h), suggesting a possibility that they may receive direct inputs from the DS-RGCs. HCR staining of *Tg(mafaa-hs:Gal4);Tg(UAS:GFP)* larvae revealed that the majority of *mafaa*+ neurons in this region are GABAergic and not glutamatergic (Fig. 5i‒m). Ca^2+^ imaging using *Tg(mafaa-hs:Gal4FF);Tg(UAS:GCaMP6s)* larvae revealed that *mafaa*-positive neurons responded exclusively to temporal motion presented to the contralateral eye (Fig. 5n,o, Extended Data Figs. 4 and 5), suggesting that these neurons correspond to a specific functional type, previously termed MoTL (<u>Mo</u>nocular <u>T</u>emporalward motion presented in the <u>L</u>eft eye) and MoTR (<u>Mo</u>nocular <u>T</u>emporalward motion presented in the <u>R</u>ight eye) (Kubo et al., 2014). Second, we labeled *nkx1.2lb*-positive neurons using a Gal4 knock-in transgenic line of *nkx1.2* gene *Tg(nkx1.2lb-hs:Gal4FF)*. *nkx1.2lb+* neurons labeled in *Tg(nkx1.2lb-hs:Gal4FF);Tg(UAS:GCaMP6s)* larvae were located in the medial part of the pretectum (Fig. 6a‒c), consistent with its endogenous expression pattern (Fig. 4f,h). Interestingly, *nkx1.2lb*-positive neurons extended commissural projections that crossed the midline through the posterior commissure (pc) and reached the contralateral pretectum (Fig. 6c). To verify that commissural projection through the pc is derived from the pretectal neurons but not from other neurons outside of the pretectum labeled in the *Tg(nkx1.2lb-hs:Gal4FF)* line, we took advantage of the pretectum-specific expression of *tcf7l2* and crossed *Tg(nkx1.2lb-hs:Gal4FF);Tg(UAS:loxP-tdTomatoCAAX-loxP-GCaMP6s)* fish with *Tg(tcf7l2-hs:Cre)* fish (Fig. 6d). This intersectional genetic strategy exclusively labels the *tcf7l2-*positive (pretectal) population of *nkx1.2lb*+ neurons. We observed that both the Gal4+, Cre- (tdTomato-positive), and Gal4+, Cre+ (GCaMP-positive) populations crossed the midline, indicating that both pretectal *nkx1.2lb+* neurons and nonpretectal *nkx1.2lb+* neurons are the source of the commissural projections (Fig. 6e–g). HCR staining of *Tg(nkx1.2lb-hs:Gal4FF);Tg(UAS:GFP)* larvae revealed that the majority of *nkx1.2lb*+ neurons are GABAergic and not glutamatergic (Fig. 6h‒l). Ca^2+^ imaging using *Tg(nkx1.2lb-hs:Gal4FF);Tg(UAS:GCaMP6s)* larvae revealed that *nkx1.2lb*-positive neurons responded broadly to the optic flow stimulus (Fig. 6m,n). They covered most of the optic flow-response types that were found in the pan-neuronal (Extended Data Fig. 2) and *Tg(tcf7l2-hs:Gal4FF)* (Fig. 3) lines, and no noticeable bias of the labeled response types was found in the *nkx1.2lb*-positive neurons. Additionally, the spatial distribution of the optic flow-responsive neurons detected from the *nkx1.2lb*-positive neurons did not differ grossly from *elavl3*- or *tcf7l2*-labeled neurons (Fig. 6n and Extended Data Fig. 4 and 5). Together, our genetic targeting of the different subtypes of optic flow-responsive cells in the pretectum revealed functional and anatomical specializations.

**Fig. 5:**
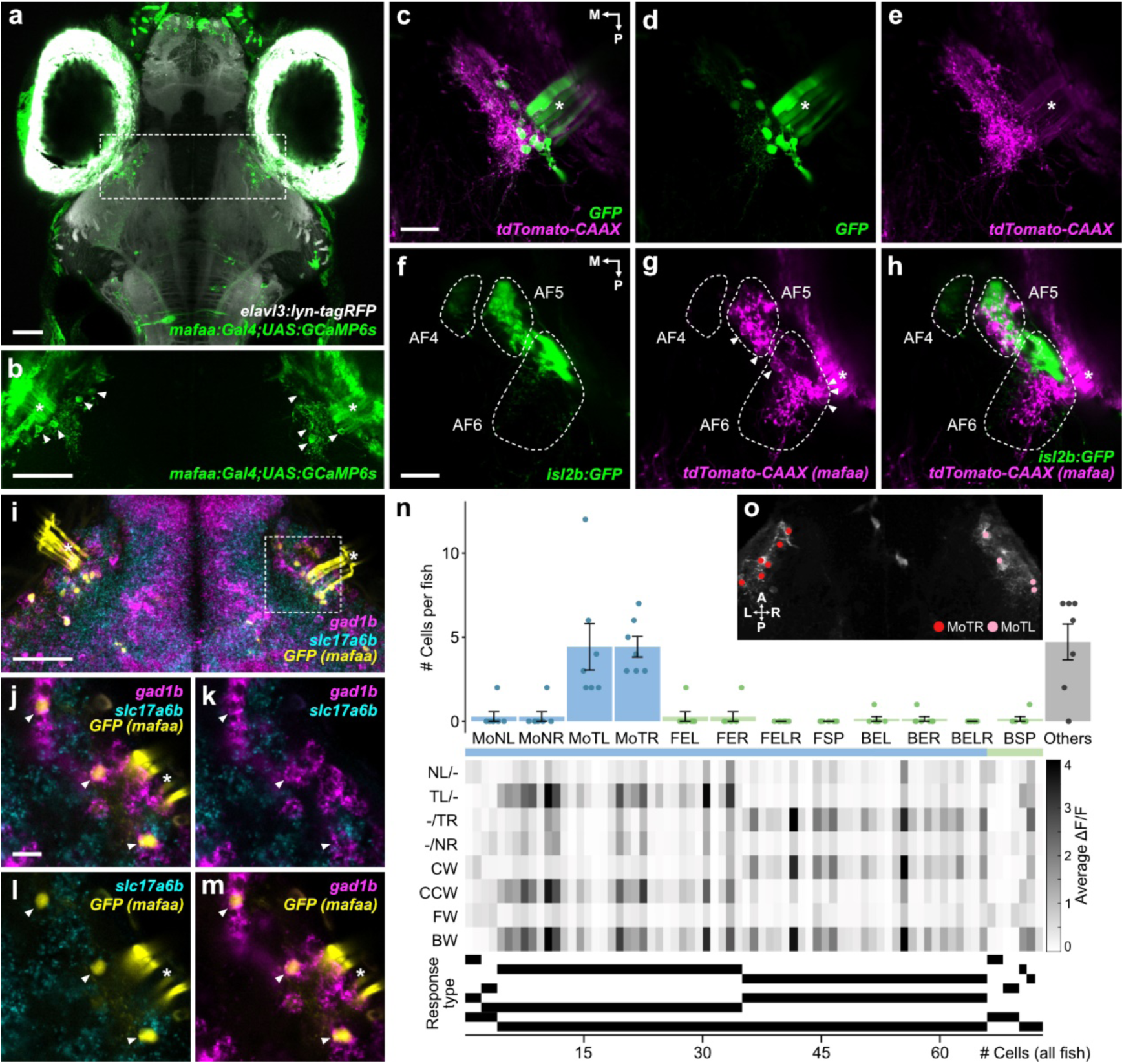
Anatomical and functional characteristics of *mafaa+* pretectal neurons. **a,b,** Substack maximum z-projections of *Tg(mafaa-hs:Gal4FF);Tg(UAS:GCaMP6s);Tg(elavl3:lyn-tagRFP)* larva. Asterisks in (**b**) denote extraocular muscles. **c-e,** Substack maximum z-projections of *Tg(mafaa-hs:Gal4FF);Tg(UAS:GFP);Tg(UAS:loxp-tdTomatoCAAX-loxp-GCaMP6s)* larva (right hemisphere). Note that membrane-targeted tdTomato-labeled neurites are locally extended in the vicinity of the soma expressing GFP. **f-h,** Distribution of *mafaa*+ neurons and the retinal ganglion cell terminals in *Tg(mafaa-hs:Gal4FF);Tg(UAS-hs:loxp-tdTomatoCAAX-loxp-GCaMP6s);Tg(isl2b:GFP)* larva. **i-m,** Substack maximum z-projections of double HCR-FISH stains of *gad1b* and *slc17a6b* in *Tg(mafaa-hs:Gal4FF);Tg(UAS:GFP)* larva (**i**). Single optical section of the region marked with dotted box in **i** (**j**-**m**). **n,** (Top) A histogram of the frequent response types in *Tg(mafaa-hs:Gal4FF);Tg(UAS:GCaMP6s)* fish (“simple”, blue, “translation-selective”, green). The remaining response types are listed as unclassified (grey). # of cells per fish represents average of 7 fish ± SEM (*n* = 7 fish). (Middle) Raster plot showing the average responses to the eight stimulus phases for all cells categorized as one of the frequent response types (*n* = 73 cells, pooled from 7 fish). Cells are ordered according to their correlation coefficient to the corresponding regressor (within each response type). (Bottom) Binary response type barcode of each cell. **o,** Spatial distribution of functionally identified cells in a representative *Tg(mafaa-hs:Gal4FF);Tg(UAS:GCaMP6s)* fish. ROIs distributed across 60 μm in depth (separated by 10 μm) are projected onto an average projection image of the GCaMP6s signals from six planes. In this volume, MoTL and MoTR response types were detected and very few of other response types were detected (for the other response types, see Extended Data Fig. 4 and 5). Arrowheads and asterisks denote *mafaa*+ neuron cell bodies and extraocular muscles, respectively. A, anterior; P, posterior; L, left; R, right; M, medial. Scale bars, 50 μm (**a**,**b**,**i**), 20 μm (**c**,**f**) and 10 μm (**j**).

**Fig. 6:**
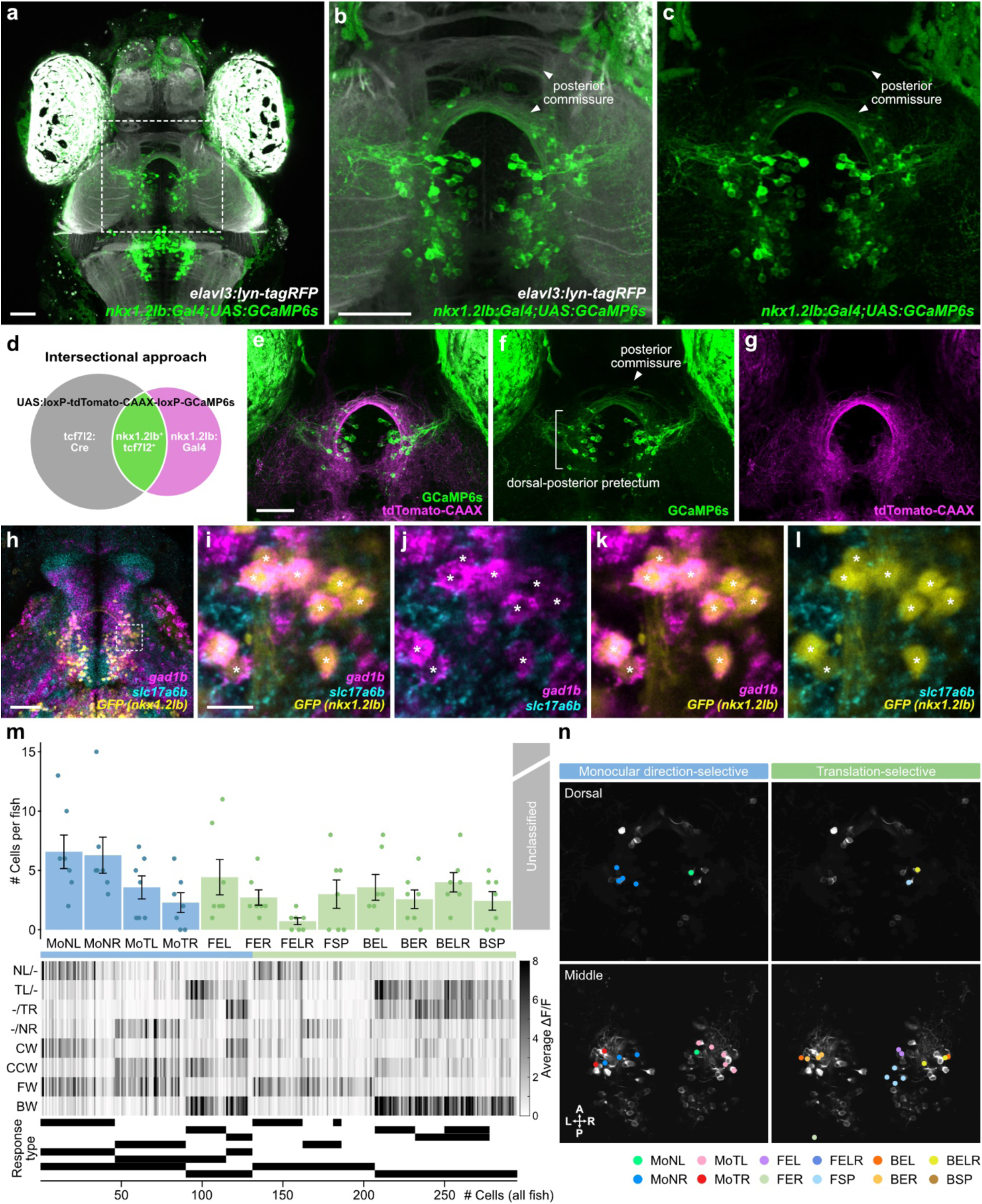
Anatomical and functional characteristics of *nkx1.2lb+* pretectal neurons. **a-c,** Substack maximum z-projections of *Tg(nkx1.2lb-hs:Gal4FF);Tg(UAS:GCaMP6s);Tg(elavl3:lyn-tagRFP)* larva (**a**) corresponds to the pretectal region marked by the dotted box in **a** (**b**-**c**). Note that GCaMP6s-labeled neurites cross the midline via the posterior commissure. **d,** Schematics for Gal4 and Cre intersectional strategy to refine genetic access to *nkx1.2lb*+ pretectal neurons. In a *nkx1.2lb Gal4* driver line, *nkx1.2lb*+ neurons activate expression of *tdTomatoCAAX* through the *UAS-hs:loxP-tdTomatoCAAX-loxP-GCaMP6s* reporter. Additional pretectal *Tg(tcf7l2-hs:Cre)* line results in *nkx1.2lb*+ pretectal neurons switching to *GCaMP6s* expression, while *nkx1.2lb*+ non-pretectal (*tcf7l2*-) cells remain to express *tdTomatoCAAX*. **e-g,** Substack maximum z-projections of a *Tg(nkx1.2lb-hs:Gal4FF);Tg(UAS-hs:loxp-tdTomatoCAAX-loxp-GCaMP6s);Tg(tcf7l2-hs:Cre)* larva. Note that GCaMP6s-labeled neurites, as well as tdTomatoCAAX-labeled neurites, cross the midline through the posterior commissure. **h-l,** Substack maximum z-projections of double HCR-FISH stains of *gad1b* and *slc17a6b* in a *Tg(nkx1.2lb-hs:Gal4FF);Tg(UAS:GFP)* larva (**h**). Single optical section of the region marked with dotted box in **h** (**i**-**l**). Asterisks denote *nkx1.2lb*+ cells that express *gad1b*. **m,** (Top) A histogram of the frequent response types in *Tg(nkx1.2lb-hs:Gal4FF);Tg(UAS:GCaMP6s)* fish (“simple”, blue, ‘‘translation-selective”, green). The remaining response types were listed as unclassified (grey). # of cells per fish represents average ± SEM of 7 fish (*n* = 7 fish). (Middle) Raster plot showing the average responses to the eight stimulus phases for all cells categorized as one of the frequent response types (*n* = 295 cells pooled from 7 fish). Cells are ordered according to their correlation coefficient to the corresponding regressor (within each response type). (Bottom) Binary response type barcode of each cell. **n,** Spatial distribution of monocular direction-selective neurons (left) and translation-selective neurons (right) from a representative *Tg(nkx1.2lb-hs:Gal4FF);Tg(UAS:GCaMP6s)* larva. Cells are color-coded according to their response type. Average image of GCaMP6s signal of the dorsal and middle representative planes of the pretectum is used as a background. For the middle plane, cells detected in the same plane, as well as those detected in 10 µm above and 10 µm below the corresponding plane, are plotted. For the dorsal plane, cells detected in the same plane, as well as those detected in 10 µm above the corresponding plane, are plotted. For the data from all individual planes, see Extended Data Fig. 4 and 5. A, anterior; P, posterior; L, left; R, right. Scale bars, 50 μm (**a**,**b**,**e**,**h**) and 10 μm (**i**).

### *nkx1.2lb*-positive neurons are required for processing the translational optic flow

Next, we tested whether the molecularly identified subtypes of the pretectal population are required for the optic flow-dependent behavior. Taking advantage of the relatively specific expression pattern of *nkx1.2b* compared to *mafaa* (which is expressed in other neurons and extraocular muscles), we selectively ablated *nkx1.2lb*+ pretectal neurons and tested the fish for two types of behavior, which are the optokinetic response (OKR) and optomotor response (OMR) (Portugues and Engert, 2009). The OKR is generally induced by clockwise or counter-clockwise rotational optic flow, while the OMR is induced by forward and backward translational optic flow. To specifically ablate the *nkx1.2lb*+ neurons located in the pretectum, we took the intersectional approach and generated *Tg(nkx1.2lb-hs:Gal4FF);Tg(UAS:loxP-TagBFP-loxP-epNTR-mScarlet);Tg(tcf7l2-hs:Cre)* triple-transgenic larvae (Fig. 7a). In these larvae, zebrafish-codon optimized enhanced-potency nitroreductase (epNTR) is specifically expressed by *nkx1.2lb*+ neurons located in the pretectum. epNTR converts the prodrug metronidazole (Mtz) into a cytotoxic compound (Curado et al., 2008; Tabor et al., 2014). The administration of Mtz to these larvae selectively and effectively ablated *nkx1.2lb*+ pretectal neurons (Fig. 7b–d). First, we tested OKR by presenting moving gratings using the visual stimulus arena that surrounds 360 degrees of the fish’s field of view (Fig. 7e). We found no significant difference in the OKR index (see Method for details) between the control and ablated fish (Fig. 7f and Extended Data Fig. 6). Second, we tested the OMR by presenting a moving grating stimulus underneath the elongated tank and recording the position of the larvae before and after the presentation of the moving grating stimuli (Fig. 7e). While groups of control larvae swam in the direction of the moving grating, groups of ablated larvae swam a significantly shorter distance (Fig. 7g,i, p = 0.03788). We found no significant effect of ablating *nkx1.2lb*+ pretectal neurons on spontaneous locomotor activity (Fig. 7h and Extended Data Fig. 6), suggesting that basal swimming activity remained intact in the ablated fish. In conclusion, commissural inhibitory *nkx1.2lb*+ neurons are required specifically for OMR but not for OKR. This suggests that *nkx1.2lb*+ neurons are required for processing translational optic flow but not rotational optic flow (see the Discussion section).

**Fig. 7:**
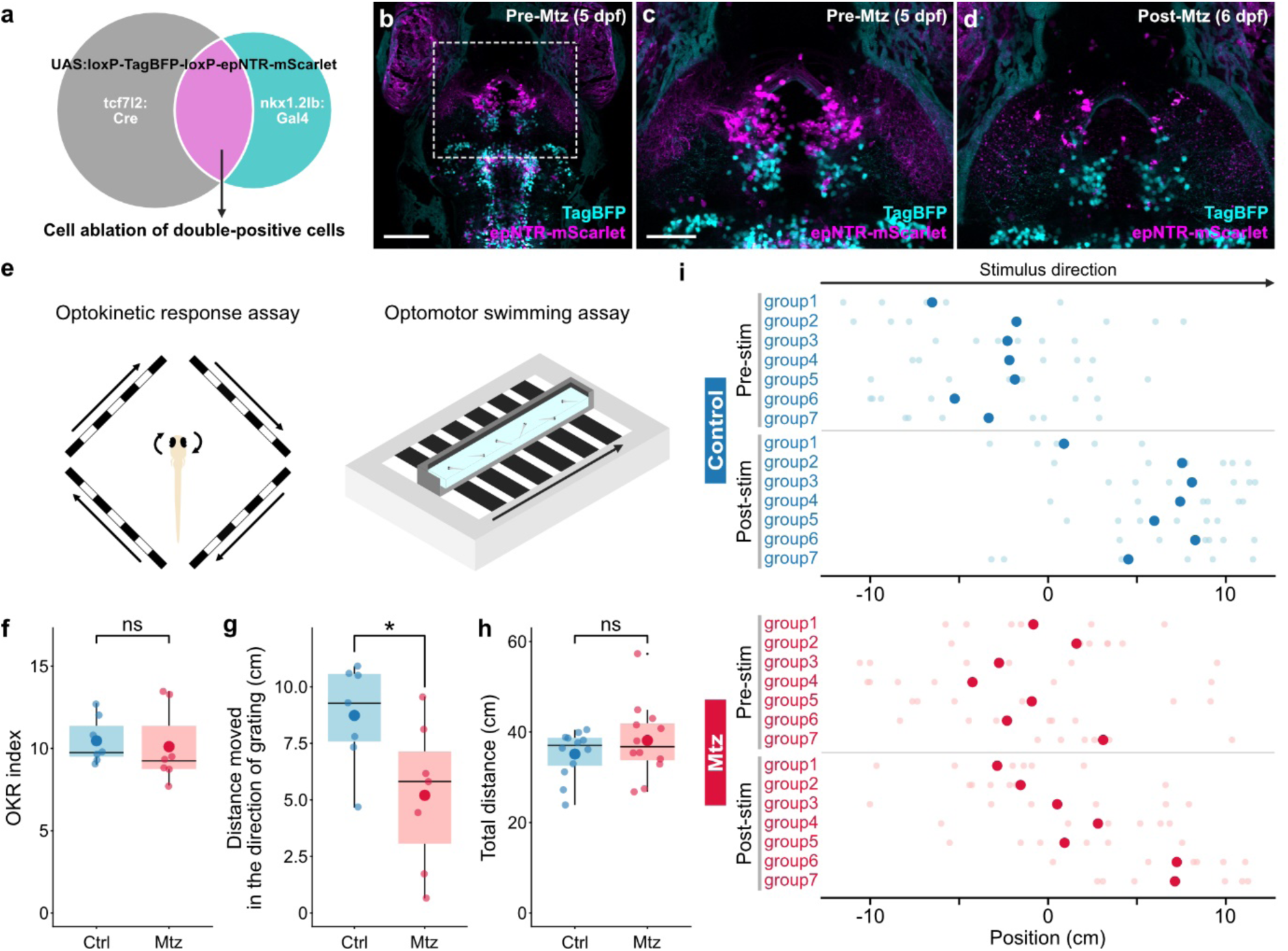
*nkx1.2lb*+ pretectal neurons are required for processing translational optic flow. **a,** Schematics for Gal4 and Cre intersectional strategy to refine genetic access to *nkx1.2lb*+ pretectal neurons. In a *nkx1.2lb* Gal4 driver line, *nkx1.2lb*+ neurons activate expression of TagBFP through a *UAS:loxP-TagBFP-loxP-epNTR-mScarlet* reporter. Additional pretectal *Tg(tcf7l2-hs:Cre)* line results in *nkx1.2lb*+ pretectal neurons switching to epNTR-mScarlet expression, while *nkx1.2lb*+ non-pretectal cells continue to express TagBFP. **b-d,** Maximum z-projections of substack of a *Tg(nkx1.2lb-hs:Gal4FF);Tg(UAS-hs:loxp-TagBFP-loxp-epNTR-mScarlet);Tg(tcf7l2-hs:Cre)* transgenic fish before (**b**,**c**) and after (**d**) treatment with metronidazole (Mtz). Scale bars, 100 μm (**b**) and 50 μm (**c**). **e,** Optokinetic response and optomotor swimming assay. **f,** Box plot showing the OKR index. The OKR index was calculated by counting the saccades in the expected direction during 1 min of grating stimulation (Mann-Whitney U test, n.s., p = 0.4047). **g,** Box plot showing the average distance moved in the direction of motion during OMR for control and ablated (Mtz) fish. Each small dot represents one group of fish and a large dot represents the mean of 7 groups (Welch two sample t-test, ∗p < 0.05, p=0.03788). **h,** Box plot showing the locomotor activity of control (*n* = 12) and ablated (Mtz, *n* = 12) fish. Total distance moved during 90 second is shown. Each small dot represents one fish and a large dot represents the mean of 12 individuals (Welch two sample t-test, n.s., p=0.3038). **i,** Position of fish before (pre-stim) and after (post-stim) the exposure to a moving visual stimulus (30 sec). Each small dot represents one of the 6 fish tested in one group and large dot indicates the mean position of the 6 fish in a group. Control larvae are either *Tg(nkx1.2lb-hs:Gal4FF);Tg(UAS:loxp-TagBFP-loxp-epNTR-mScarlet);Tg(tcf7l2-hs:Cre)* transgenic larvae untreated with Mtz or *Tg(nkx1.2lb-hs:Gal4FF);Tg(UAS:loxp-TagBFP-loxp-epNTR-mScarlet)* transgenic larvae treated with Mtz.

## Discussion

By strategically combining the functional labeling of neurons using CaMPARI and subsequent scRNA-seq, we developed a technique to analyze the transcriptomic features of functionally active neurons during optic flow stimulation in larval zebrafish. Most pretectal neurons expressed the *tcf7l2* gene and encompassed most of the previously characterized optic flow response types. Further clustering revealed several transcriptomic subtypes of pretectal neurons that were distinguishable from one another by anatomical and functional characteristics. Finally, we molecularly identified inhibitory commissural pretectal neurons that were necessary for optic flow-dependent behavior, specifically for the OMR, which depends on the translational optic flow; however, it is dispensable for the OKR, which depends on the rotational optic flow.

In this study, we integrated the functional tagging of neurons by CaMPARI2 and subsequent single-cell transcriptomic analyses. Although the physiological properties and transcriptomic profiles of single cells have generally been independently studied, a couple of methods have been described to link the physiological and transcriptomic features of individual neurons (Yagishita and Sasaki, 2024). One such method, Patch-seq, is based on the electrophysiological recording and subsequent RNA-seq analysis of single neurons (Cadwell et al., 2016; Fuzik et al., 2016; Scala et al., 2021). Although it is a powerful tool to study both aspects of the neuronal diversity of the same cells, the throughput of cell sampling is generally limited. In contrast, the optical tagging of functional neuron types and the subsequent RNA-seq analysis we employed in this study offer the advantage of sampling thousands of neurons simultaneously. Furthermore, since CaMPARI2-based labeling uses light, it provides a precise temporal window for labeling neurons of interest, compared with other techniques based on immediate early gene promoters, which have been primarily used for behavior where the temporal kinetics are relatively slow (Chen et al., 2020; Sun et al., 2024). A combination of CaMPARI2 and scRNA-seq applied in the mouse identified previously known transcriptomic types of layer 2/3 neurons in the visual cortex and validated that these transcriptomic types exhibit distinguishable negative and positive prediction error responses (O’Toole et al., 2023). Our current study extends this research by identifying previously unknown molecular markers for the behaviorally relevant functional cell types in the pretectum.

A previous scRNA-seq study characterized the transcriptomic diversity of neurons in the diencephalon, including the pretectum, thalamus, and prethalamus, as well as several surrounding brain regions in the larval zebrafish (Sherman et al., 2023). Some of the pretectal marker genes identified in their dataset, such as *npy*, *penkb*, *and zic1*, were also detected in our dataset. Most of these clusters are consistently GABAergic in our data and that of Sherman et al.’. The *mafaa* and *nkx1.2lb* genes that we focused on in our study were not described as representative marker genes in the study by Sherman et al. We speculate that our functionally targeted approach successfully dissected out a rare and functionally relevant population of neurons that have been missed in the conventional anatomically based approach.

Binocular integration of the optic flow critically depends on the interhemispheric transfer of DS information in the pretectum (Kubo et al., 2014; Naumann et al., 2016). Previous anatomical studies conducted using single-neuron reconstructions and tracer experiments in larval and adult zebrafish pretectum have demonstrated that some pretectal neurons cross the brain side through either of the two commissural fibers: the posterior commissure (pc) or postoptic commissure (poc) (Baier and Wullimann, 2021; Kramer et al., 2019; Yáñez et al., 2018). The pc represents a conserved landmark located in the dorsal part of the pretectum of vertebrates, while the poc is located at the extreme ventral part of the diencephalon (Wilson et al., 1990). Furthermore, electron microscopy-based reconstruction of functionally identified optic flow-responsive cells in larval zebrafish revealed that the optic flow-responsive cells frequently cross the midline via either the pc or poc and project to the contralateral pretectum (Svara et al., 2022). Our study identified *nkx1.2lb* as a specific marker for pretectal neurons that cross the midline via the pc. Genetic markers for the poc-crossing pretectal neurons are yet to be identified. Contralaterally projecting pretectal neurons identified in the EM reconstruction had dendrites in ipsilateral retinorecipient neuropil areas that receive DS inputs (AF5 and AF6) and/or medial pretectal neuropil (mPN) and projected axons to the contralateral mPN and/or AF6 in the pretectum (Svara et al., 2022). The similarity between their anatomical features suggests that *nkx1.2lb*+ commissure neurons may receive direct DS retinal inputs and innervate pretectal neurons located on the contralateral side (Fig. 8). Future studies are needed to identify the pre- and postsynaptic neurons of the *nkx1.2lb*+ pretectal neurons.

**Fig. 8:**
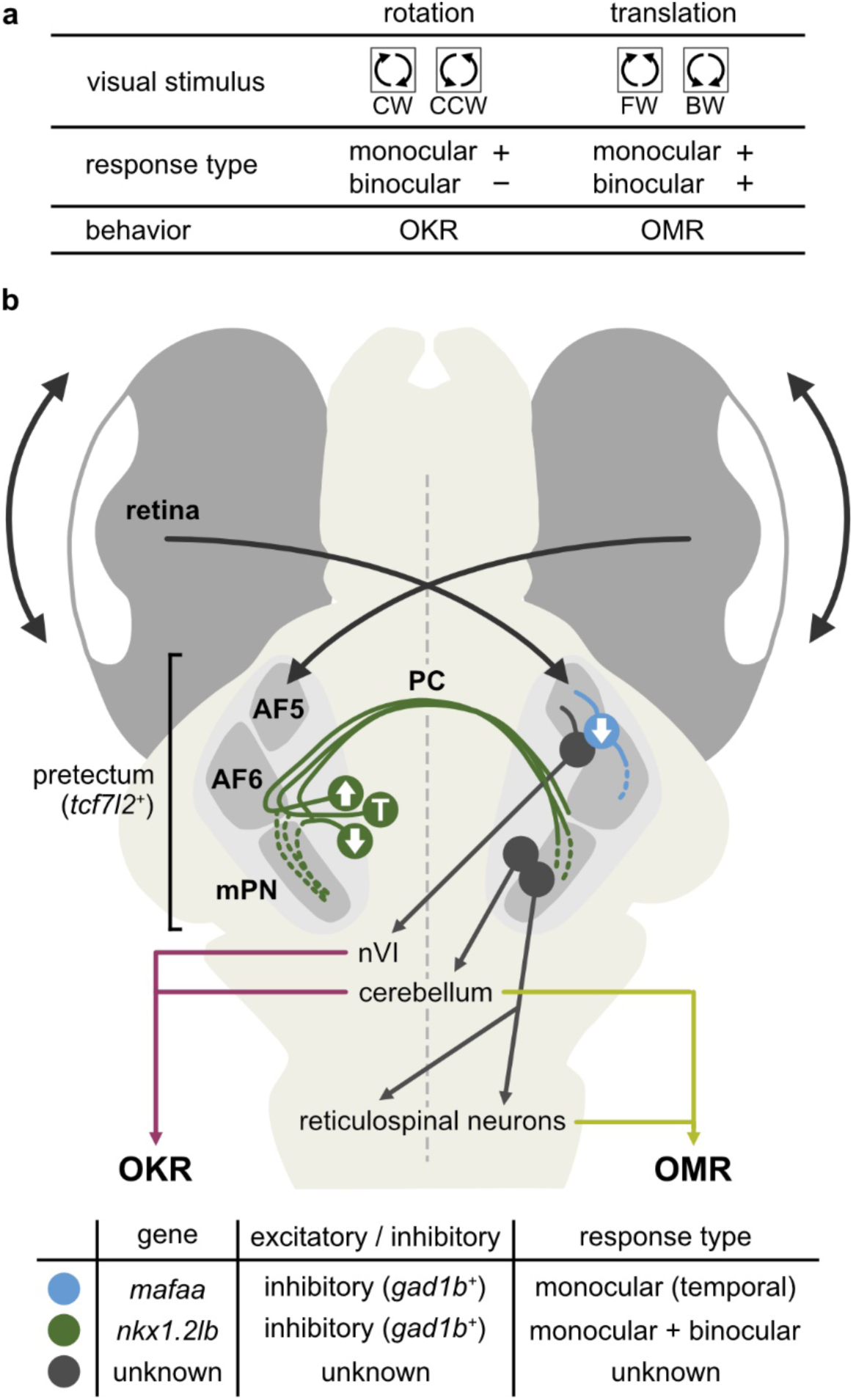
Model for the optic flow processing circuit for controlling OKR and OMR. **a,** Summary of previous results (Kubo et al., 2014). Clockwise (CW) and counter-clockwise (CCW) rotational optic flow mainly induces the OKR. In the pretectum, monocular optic flow-responsive cells are active during rotational optic flow, whereas rotation-selective binocular cells are rarely observed. For processing of the forward (FW) and backward (BW) translational optic flow that induces OMR, monocular optic flow-responsive cells and binocular translation-selective cells are active. **b,** Proposed circuit model. Pretectal output neurons are depicted based on 1) projection neurons that project to cerebellum and reticular formation in the hindbrain (Kramer et al., 2019) and 2) pretectal neurons that extend dendrites into AF5 (which contains the axon terminals of direction-selective RGCs) and project axons to the contralateral abducens nucleus (nVI) (Dowell et al., 2025). PC, posterior commissure; AF5, arborization field 5; AF6, arborization field 6; mPN, medial pretectal neuropil.

Our previous functional imaging study proposed a parsimonious circuit wiring diagram of the optic flow processing circuit in the pretectum, where the monocular and binocular response types are hierarchically organized to compute the rotational and translational optic flow (Kubo et al., 2014). The current study provides updates on this model. First, while the previous model predicted that all of the commissural pretectal neurons would be excitatory, we found that at least some of the commissural pretectal connections are inhibitory, since the inhibitory *nkx1.2lb*+ pretectal neurons project contralaterally. The discrepancy can be explained, at least in part, by the direct or indirect connectivity of inhibitory inputs. The previous model proposed that interhemispheric inhibition is mediated indirectly by commissural excitatory relay neurons, which, in turn, activate local inhibitory neurons. We now propose that the inhibition is provided directly by commissural inhibitory neurons. Second, the previous model predicted that the monocular response types consist of both excitatory and inhibitory neurons, while the binocular response types consist exclusively of excitatory neurons. Our data suggest that the majority of pretectal transcriptomic types, including *mafaa*+ and *nkx1.2lb*+ neurons, express the inhibitory neuron maker *gad1b*. For instance, we found some of the binocular neurons to be inhibitory using the *nkx1.2lb* transgenic line. Although we do not rule out the possible contribution of excitatory neurons, the majority of optic flow-responsive neurons are inhibitory. The excitatory connections predicted in the previous model could be derived from the DS-RGC inputs, which are glutamatergic and excitatory. Together, our current study provides direct experimental evidence for anatomical projections and the sign of connectivity (i.e., excitatory vs. inhibitory) of the functional circuit model of the pretectum.

Our data showed that *nkx1.2lb*+ neurons consist of both monocular and binocular types and the ablation of pretectal *nkx1.2lb*+ neurons impaired the OMR but left the OKR intact. Prior functional imaging studies have shown that the majority of active neuron types during rotational motion in the horizontal plane are monocular simple neurons (namely MoNL, MoNR, MoTL, and MoTR), suggesting that rotational motion and the resulting rotational OKR are largely computed by monocular simple cells in the pretectum (Fig. 8) (Kubo et al., 2014; Wang et al., 2019; Zhang et al., 2022). These observations led to the idea that during rotational motion, the optic flow information from both eyes is separately processed and not combined in the pretectum (Kubo et al., 2014; Wang et al., 2019; Zhang et al., 2022). In contrast, during translational motion, binocular translation-selective cells are active in addition to the simple monocular neurons (Kubo et al., 2014; Wang et al., 2019; Zhang et al., 2022). Moreover, the activities of these binocular translation-selective cells are selectively suppressed during rotational motion, meaning that they are, either directly or indirectly, inhibited by the simple cells that respond to the motion sensed by the other eye during rotation (e.g., the FEL is inhibited by MoTR during CW motion). The intact OKR after pretectal *nkx1.2lb*+ neuron ablation suggests that the computation of rotational optic flow does not require monocular cells labeled in the *nkx1.2lb* transgenic line. This indicates that other monocular cells that were unlabeled in the *nkx1.2lb* line play essential roles in the OKR. We speculate that one such candidate is *mafaa*+ neurons located in the migrated pretectal region M1 (Baier and Wullimann, 2021; Mueller and Wullimann, 2002). Neurons in the M1 have been shown to play critical roles in OKR (Wu et al., 2020). *mafaa*+ neurons closely resemble those neurons in terms of 1) direction tuning to motion, 2) proximity to RGC arborization fields AF5 and AF6 that harbor DS-RGC axon terminals (Kramer et al., 2019), and 3) their neurite projections in the local pretectal neuropil (Wu et al., 2020). Unfortunately, pretectal *mafaa*+ neuron ablation is technically difficult at present even when using the intersectional genetics with *Tg*(*mafaa-hs:Gal4FF)* and *Tg(tcf7l2-hs:Cre)* lines, since both transgenic lines label the eye and tail muscles (see for example Fig. 2g,h and 5b,c). Given the critical role of M1 neurons in the OKR, we speculate that *mafaa*+ neurons may contribute significantly to the motion processing and/or generation of the OKR. On the other hand, the processing of the translational optic flow during OMR requires *nkx1.2lb*+ pretectal neurons. The observed OMR defect can be caused by 1) the ablation of monocular neurons that provide inhibitory inputs to generate binocular neurons, 2) the ablation of binocular neurons that encode translation-selective information, or a combination of both processes. In any case, the essential role of *nkx1.2lb*+ inhibitory neurons for OMR but not for OKR is consistent with the previous model in which processing of the rotational optic flow does not require binocular inhibitory integration while the processing of the translational optic flow and behavior requires inhibitory connectivity between hemispheres.

The cell-type-specific transcriptome is generally used to classify neuron diversity (Zeng, 2022; Zeng and Sanes, 2017) and is believed to give rise to distinct morphology, connectivity, and function (Goetz et al., 2022; Huang et al., 2022; Kölsch et al., 2021). Consistent with this view, we found that one pretectal transcriptomic type corresponds to one morphological type: *mafaa*+ and *nkx1.2lb+* pretectal neurons project locally and contralaterally, respectively. However, in terms of function, we found that *nkx1.2lb*+ neurons consisted of multiple optic flow-response types, suggesting that a single transcriptomic type can consist of multiple functional types. This supports the conclusion of a prior work showing that transcriptionally similar optic tectum neurons can be functionally divergent (Shainer et al., 2025). The functional diversity of *nkx1.2lb*+ neurons may arise from different inputs received by each neuron (e.g., nasally tuned vs. temporally tuned DS inputs).

In summary, our functionally targeted transcriptional profiling provided a molecular handle on the optic flow processing circuit and gained new insights into the circuit mechanism for computing behaviorally relevant visual information. The molecular markers we identified may help elucidate how diverse pretectal neuron types are generated and wired together to form the functional optic flow processing circuit during development. Future studies will identify comprehensive catalogs of the transcriptomic, functional, and morphological neuron types underlying the optic flow processing and synaptic connectivity of the identified neurons.

## Acknowledgements

We thank Shin-ichi Higashijima (National Institute for Basic Biology) for providing the donor plasmids for the CRISPR-Cas9 knock-in, Shachar Sherman (Max Planck Institute for Biological Intelligence), Grigorios Oikonomou, and David Prober (Caltech) for sharing cell dissociation protocols prior to publication, Shigehiro Kuraku (National Institute of Genetics) for advice on bioinformatics analysis. We thank Masahiko Hibi, Hisaya Kakinuma, Ayjan Urazbayeva, and Hitoshi Okamoto for their discussion and feedback on our manuscript. We thank the members of the Kubo laboratory for their technical assistance and advice and the Research Resource Division of RIKEN CBS for fish care and technical assistance. Bioinformatics computations (Cell Ranger analysis) were performed on the NIG supercomputer at ROIS National Institute of Genetics. We are grateful to National BioResource Project (NBRP) Zebrafish Japan for providing *Tg(UAS:GFP)* and *Tg(elavl3:LOXP-DsRed-LOXP-EGFP)* lines. This study was supported by JSPS Grant-in-Aid for Scientific Research (KAKENHI) (19K23787, 20K15906, 23K14293 to KM; 24K02008 to KK; 17K20147, 21H02586, 22K21353 to FK), “Strategic Research Projects” grant from ROIS (Research Organization of Information and Systems) (to KM), and Tomizawa Jun-ichi & Keiko Fund of the Molecular Biology Society of Japan for Young Scientist (to FK).

## Author contributions

KM and FK conceived and designed the study, and wrote the paper with inputs from CHW, AT, TS and KK. KM, CHW, AT and FK conducted experiments. KM analyzed the data. TS and KK generated the pTol2 UAS-hsp:epNTR-mScarlet plasmid.

## Declaration of interests

Authors declare no competing interests.

## Methods

### CONTACT FOR REAGENT AND RESOURCE SHARING

Further information and requests for resources and reagents should be directed to the Lead Contact Fumi Kubo.

### EXPERIMENTAL MODEL AND SUBJECT DETAILS

#### Animal care and transgenic zebrafish

Adult and larval zebrafish (*Danio rerio*) were housed and handled according to standard procedures (Westerfield, 2007). Animal experiments were performed according to regulations of National Institute of Genetics and RIKEN Center for Brain Science. We used the following previously described transgenic lines: *Tg(UAS:GFP)nns19, Tg(UAS:GCaMP6s)mpn156, Tg(UAS:mCherry)s1984t, Tg(elavl3:LOXP-DsRed-LOXP-EGFP)nns16Tg, Tg(elavl3:H2B-GCaMP6s)jf5Tg, Tg(elavl3:lyn-tagRFP)mpn404* (a.k.a. HuC:lyn-tagRFP). Transgenic fish were kept in either a TL or TLN (nacre) background and larvae lacking trunk pigmentation (outcrossed to TLN, nacre) were used in the experiment. Zebrafish larvae were raised in E3 solution on a 14/10 h light/dark cycle until 5 or 7 days post-fertilization (dpf).

#### Transgenic line establishment

To generate the pTol2-elavl3*:NLS-CaMPARI2* plasmid, the CaMPARI2 coding sequence was amplified from pAAV_hsyn_NES-his-CaMPARI2-WPRE-SV40 plasmid (gift from Dr. Erick Schreiter, Addgene plasmid #101060), fused with nuclear localization signal (NLS), and inserted into a pTol2 vector that contains elavl3 pan-neuronal promoter. To generate the pTol2-UAS-hsp:epNTR-mScarlet plasmid, fragments containing heatshock promoter (hsp) sequence from the UAShspzGCaMP6s (Muto et al., 2017), epNTR (codon optimized for zebrafish, Tabor et al., 2014) and mScarlet (codon optimized using CodonZ, Horstick et al., 2015) were inserted between UAS and SV40 sequences synthesized pTol2-UAS vector using In-Fusion cloning. To generate the pTol2-UAS-hs:loxP-tdTomatoCAAX-loxP-GCaMP6s plasmid, three fragments for loxP-tdTomatoCAAX (Förster et al., 2017), SV40-loxP and GCaMP6s (Thiele et al., 2014) were PCR amplified and cloned between a hsp sequence and SV40 sequence of pTol2-UAS-hsp:epNTR-mScarlet plasmid by In-Fusion cloning (Takara). To generate the pTol2-UAS-hsp:loxP-TagBFP-loxP-epNTR-mScarlet plasmid, epNTR-mScarlet were inserted into the UAS-hsp:loxP-TagBFP-loxP vector. *Tg(elavl3:NLS-CaMPARI2), Tg(UAS-hs:loxP-tdTomatoCAAX-loxP-GCaMP6s)* and *Tg(UAS-hs:loxP-TagBFP-loxP-epNTR-mScarlet)* transgenic lines were generated using the standard Tol2 transposon system.

*Tg(tcf7l2-hs:Gal4FF)*, *Tg(tcf7l2-hs:Cre)*, *Tg(tal2-hs:Gal4FF)*, *Tg(mafaa-hs:Gal4FF)*, *Tg(nkx1.2lb-hs:Gal4FF)* lines were generated using the published method (Kimura et al., 2014) with some modifications. Briefly, single guide RNA sites were designed in a region 200-600 bp upstream of the transcription start site of the target gene using CCTop (Stemmer et al., 2015). CRISPR-Cas9 RNP complex was prepared according to the manufacturer’s instructions (IDT). In short, gRNA was produced by annealing custom-synthesized crRNA (Alt-R CRISPR-Cas9 crRNA, IDT) with tracrRNA (IDT, CAT# 1072532) in Nuclease-Free Duplex Buffer (IDT, CAT# 11-05-01-12) and subsequently incubated with Cas9 protein (IDT, CAT# 1081060) at 37°C for 15 minutes. Another CRISPR-Cas9 RNP complex targeting either G-bait or M-bait sequence of the donor plasmid was prepared separately. Donor plasmid containing Gbait-hsp70:Gal4FF, Mbait-hsp70:Gal4FF or Mbait-hsp70:Cre (Kimura et al., 2014) (gift from Dr. Shin-ichi Higashijima) was added to a final concentration of 25 ng/μl. The freshly prepared mix of two CRISPR-Cas9 RNP complexes and the donor plasmid was injected into the cell of *Tg(UAS:mCherry)*, *Tg(UAS-hs:loxP-tdTomatoCAAX-loxP-GCaMP6s)*, *Tg(UAS:GCaMP6s)* or *Tg(elavl3:LOXP-DsRed-LOXP-EGFP)* transgenic zygotes. Transient expressors were raised and screened for germline transmission. Gal4- or Cre-positive larvae (F1 fish) were used to confirm correct insertion of the donor plasmid into the target gene locus by PCR using primer pairs designed to amplify fragments that span the region between the genomic locus of the target gene and the donor plasmid.

#### CaMPARI photoconversion during visual stimulation

To perform photoconversion of CaMPARI2, we used a custom-built microscope equipped with a 405 nm UV LED (Thorlabs, #M405L4) and a 20x water immersion objective lens (Olympus, UMPLFLN20XW, NA0.5). The UV intensity measured at the sample was around 2.5 mW and the power density was around 326.1 mW/cm^2^. To validate the photoconversion efficacy of NLS-tagged CaMPARI2, *Tg(elavl3:NLS-CaMPARI2)* transgenic larvae between 6 and 7 dpf were embedded in 2% agarose in the center of a dish with a diameter of 3 cm and UV light was applied to the center of the pretectal area for 10 min in the presence of 800 μM 4-aminopyridine (4-AP). To photoconvert neurons during visual stimulation, *Tg(elavl3:NLS-CaMPARI2)* transgenic larvae were similarly embedded in agarose. Sinusoidal grating stimuli were generated by a custom-written script using PsychoPy and presented to a custom-built oLED arena that consists of four oLED screens (3.81 inch, Wisecoco) that covered 360 degrees around the fish. Binocular gratings moved horizontally in four phases in the following order: clockwise, counter-clockwise, forward and backward (duration: 10 s each at spatial frequency of 0.033 cycles/degree and temporal frequency of 2 cycles/sec, interspersed with 2 s stationary gratings, Extended Data Fig. 1a) with 10 times repeats. The total length of the visual stimulus protocol was about 8 min 20 sec. In separate experiments using fish embedded with eyes free, we made sure that fish showed normal optokinetic response under this UV photostimulation condition, confirming that the UV light does not interfere with the detection of the visual stimuli (data not shown). In the control visual stimulation, the same photostimulation protocol was used except that the speed of grating stimulus was set to zero (stationary gratings). Confocal imaging was performed on a Zeiss LSM900 microscope to visualize CaMPARI2 green and red fluorescent proteins. When comparing photoconversion efficiency among different conditions (Fig. 1 and Extended Data Fig. 1), same confocal imaging settings (laser power and gain) were used across conditions.

#### Cell dissociation and Fluorescence-Activated Cell Sorting (FACS)

Photoconversion of CaMPARI2 from green to red state is irreversible (Fosque et al., 2015). Consistent with this, we observed that the CaMPARI2 Red signal persisted for several hours after photoconversion. After photoconversion, larvae were kept at 27.5 degrees for maximally 4 hours until brain diessection. Brain was dissected in ice-cold Neurobasal medium (ThermoFisher Scientific, 21103049) supplemented with B-27 (ThermoFisher Scientific, 17504044). After the eyes were removed from the head, anterior to the habenula and posterior to the otolith was cut away to increase the proportion of pretectal cells (Fig. 1a). Cell dissociation was performed as previously described with some modifications (Sherman et al., 2023). Tissues were pooled from 50-60 larvae and digested in 25 U/ml papain, 130 U/ml DNaseI, 2 mM L-cysteine in oxigenated Ames at 37°C for 36 minutes. Every 12 minutes, tissues were triturated by pipetting, followed by a final incubation for 5 minutes. The cell suspension was then passed through a pre-wet 40 µm filter (Falcon, 352340). To stop the digestion, the papain solution was replaced by papain inhibitor solution containing 15 mg/ml ovomucoid and 15 mg/ml BSA. Tissue was gently dissociated by trituration using a broad end tip pipette in the papain inhibitor solution. To wash the cell suspension, cells were pelleted at 300 g for 7 minutes at RT and resuspended in FACS buffer (oxigenated Ames containing 0.25% BSA (Sigma, A4161), 10m M HEPES (Sigma, H0887), 25 U/mL DNase I (Thermofisher, 90083), 5mM MgCl_2_)).

CaMPARI2 red cells were sorted based on green and red fluorescence using SONY Cell Sorter SH800S equipped with 488 nm & 561 nm lasers. In a pilot experiment, *Tg(elavl3:NLS-CaMPARI2)* larvae without photoconversion were used to determine sorting gates. First, FACS gates were set after 50,000 recorded events to sort live single-cells. Similar gates were used across experiments. Specifically, a scatter gate was initially set using forward scattering (FSC-A) and back scattering (BSC-A) to select the live cell population. Next, Forward Scatter Area (FSC-A) and Forward Scatter Height (FSC-H) were used to eliminate debris as much as possible and isolate single-cells. Then, CaMPARI2 green and CaMPARI2 red fluorescence were used to collect CaMPARI2 red cells labeled by photoconversion of *Tg(elavl3:NLS-CaMPARI2)* larvae. The gating was set to include the CaMPARI2 red fluorescent cell population, which was not observed in the *Tg(elavl3:NLS-CaMPARI2)* control larvae that did not undergo photoconversion. Overall, 60-70% of the population consisted of live single cells, and approximately 8% of those were CaMPARI2 red cells. These cells were sorted using a 100 µm nozzle (20 PSI) and collected to a 2 ml protein LoBind Eppendorf tube filled with 1 ml oxigenated Ames containing 1.0% BSA, 10mM HEPES. A total of 130,000∼150,000 CaMPARI2 red cells were collected from 50-60 larvae. After FACS was completed, cell suspension was centrifuged for 5 min at 100 g at RT and the pellet was resuspended in 300 µl of Ames medium/0.5% BSA. This washing step was repeated three times, and finally, the cells were resuspended in 30 μL of Ames/BSA solution. To check cell density and viability, cells were stained with a dead cell stain Trypan Blue (Sigma-Aldrich, T8154) and both live and dead cells were counted using a DIC microscope. Cell density was typically between 300 and 800 cells/µl with a viability rate of over 90%.

#### Single Cell RNA-seq

Droplet RNA sequencing was performed according to the manufacturer’s instructions with no modifications (10x Genomics Chromium System, Chromium Next GEM Single Cell 3’ Reagent Kits v3.1). A single-chromium chip channel was loaded with a cell suspension, aiming to capture 5,000 cells with a doublet rate of <5%. For the “moving” condition, 2 replicates were collected from 2 experiments. For the “stationary” condition, 2 replicates were collected from 2 experiments. cDNA libraries were sequenced on a NovaSeq 6000 (illumina) or DNBSEQ-G400 (MGI Tech) to a depth of ∼60,000 reads per cell.

#### Computational analysis of single cell transcriptomics data

##### Alignment of gene expression reads, preprocessing and batch integration

Initial preprocessing was performed using the Cell Ranger software suite (version v6.0.1, 10X Genomics) following standard guidelines. ‘‘cellranger count’’ was used to align the reads to a custom reference genome created by combining the improved zebrafish transcriptome annotation file (GTF for v4.3.2) (Lawson et al., 2020) and the FATSA file (GRCz11) with CaMPARI2, using ‘‘cellranger mkref’’. Each biological replicate resulted in 3,000-5,000 estimated number of cells with 3,500-5,000 median unique molecular identifier (UMI) counts per cell. Output from cellranger was loaded into Seurat R package (Satija et al., 2015) (version 4.1.1) using the Read10X function, with each condition representing different experiment: stationary1, stationary2, moving1, and moving2. The data were stored as Seurat objects, with a minimum threshold of 200 genes per cell and a minimum of 3 cells per gene for inclusion. For quality control, cells with greater than 10% mitochondrial gene expression were excluded from further analysis. Cells were also filtered based on the number of detected genes, retaining those with between 200 and 5000 genes. This quality control was applied uniformly to all datasets. Following filtering, the datasets were integrated using the SCTransform function. The datasets for stationary1, stationary2, moving1, and moving2 were stored in a list and processed through the SCTransform function. For each transformed dataset, the 3000 most variable genes were selected as possible integration features using SelectIntegrationFeatures. We then followed the standard integration pipeline, employing PrepSCTIntegration to prepare the data for integration. Integration was performed using the FindIntegrationAnchors function, where stationary1 and stationary2 datasets were used as the reference. Normalization was done using the SCTransform. The resulting integration anchors were used to combine the datasets into a single Seurat object using IntegrateData. To reduce dimensionality, principal component analysis (PCA) was performed using the RunPCA function. Subsequently, UMAP was generated using the first 20 principal components (PCs) to visualize the data in a two dimensional space (RunUMAP). Clustering was performed using the FindNeighbors and FindClusters functions with the PCA reduction and a resolution of 0.8. Marker genes for each cluster were identified using the FindAllMarkers function, selecting only positive markers with a log fold change threshold of 0.25. We annotated clusters based on the expression data of marker genes using the Zebrafish Information Network website (zfin.org).

##### Identification of optic flow-responsive pretectal cluster

To identify clusters in which optic flow responsive cells are enriched, we calculated the composition of different clusters within each dataset and compared them across moving and stationary conditions. For each cluster, we calculated the fold change of the proportion of “moving” datasets (average of the two replicates) with respect to that of “stationary” datasets (average of the two replicates) (MS index) using the following formula

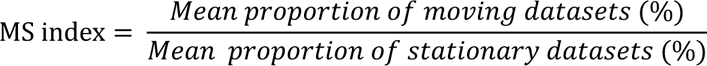

Among the 16 clusters that showed MS index higher than 1 (Fig. 1f), we excluded 8 clusters since their identities were unambiguously annotated based on published data (Extended Data Fig. 1h). Cluster 7 was classified as telencephalon/prethalamus due to the expression of *dlx5a* and *arxa*, cluster 9 as hindbrain based on the expression of *slc30a2* and *hoxb3a*, cluster 11 as dorsal habenula due to the expression of *gng8* and *GPR151*, cluster 16 as trigeminal sensory neurons based on *p2rx3b* and *p2rx2*, cluster 17 as ventral habenula due to the expression of *kiss1* and *GPR139*, cluster 22 as cerebellum based on the expression of *gsg1l* and *trpc3*, and cluster 26 as hypothalamus based on the expression of *pdyn* and *oprd1b*. Cluster 23, which expressed of *krt4* and *krt91*, was determined not to be neuronal. The remaining 8 clusters, “2”, “3”, “6”, “8”, “12”, “15”, “20”, and “21” were considered as candidate clusters that potentially contain optic flow responsive cells in the pretectum (Fig. 1f). The subset was split by the “orig.ident” variable, and 2000 variable genes were identified for each subset using the “vst” selection method. The datasets were integrated using SelectIntegrationFeatures and FindIntegrationAnchors, followed by integration using the IntegrateData function. The standard workflow for clustering and visualization was performed, including normalization, PCA (with 10 PCs), and UMAP. Clustering was performed with a resolution of 0.8. Marker genes for each cluster were identified using the FindAllMarkers function, selecting only positive markers with a log fold change threshold of 0.25. As a result, 16 new clusters were obtained (Fig. 2), and the MS index calculation and brain region annotation were performed as described above. Re-clustering of the pretectum cluster (cluster 5) was performed in a similar manner (Fig. 4). However, due to the low number of cells, the k.weight parameter in the IntegrateData function was set to the minimum sample size of 68 (moving1: 165 cells, moving2: 132 cells, stationary1: 68 cells, stationary2: 106 cells).

#### Immunohistochemistry

Larvae (6 dpf) were fixed in 4% PFA/PBS at 4°C overnight and processed for the whole-mount antibody staining. Samples were incubated with primary antibodies for 7 days at 4°C and then with secondary antibodies for 5 days at 4°C. Primary antibodies used are as follows: chicken anti-GFP (1:1000, Invitrogen, A10262), mouse anti-TCF4 (also known as TCF7L2) (1:400, 6H5-3, Millipore), rabbit anti-Synapsin 1/2 (1:1000, Synaptic Systems, #106002). Anti-chicken Alexa 488 (1:250, ThermoFisher, A-11039), anti-mouse Alexa 555 (1:250, ThermoFisher, A-21127) and anti-rabbit Alexa 647 (1:250, ThermoFisher, A-21244) were used as secondary antibodies. Confocal imaging was performed on a Zeiss LSM900 microscope.

#### Hybridization chain reaction (HCR) fluorescent in situ RNA labeling (HCR-FISH)

Larvae (5-6 dpf) were fixed in 4% PFA at 4°C overnight and processed for HCR fluorescent in situ RNA labeling (HCR-FISH) according to the manufacturer’s instructions (Molecular Instruments). *Tg(tcf7l2-hs:Gal4FF);Tg(UAS:GFP), Tg(mafaa-hs:Gal4FF);Tg(UAS:GFP)*, *Tg(nkx1.2lb-hs:Gal4FF);Tg(UAS:GFP)* larvae were hybridized with two probe sets, each of which was detected with Alexa546 or Alexa647 amplifiers. Imaging was performed on a Zeiss confocal microscope (LSM900).

#### Functional Calcium imaging

Larvae expressing GCaMP were mounted in agarose (LMP-agarose, 2% w/v in E3 medium). We used a two-photon microscope (BergamoII, Thorlabs) equipped with a 20x water immersion objective (XLUMPLFLN 20x, NA1.0, Olympus) and a fiber laser with a fixed-wavelength at 920 nm (ALCOR-920, Spark Lasers). The visual stimuli were presented to the fish using a custom-built oLED arena that consists of four flat oLED screens (3.81 inch, Wisecoco) arranged in square and covered 360 degrees around the fish. To ensure monocular stimulation, the central 45° portion of the visual field was covered with black paper. Sinusoidal grating stimuli were generated by a custom-written script using PsychoPy. RGB color space of the oLED screens were set to [1.0, -1, -1] (red stimulus). Furthermore, magenta gelatin filter (No. 32, Kodak) was applied in front of the oLED screens to block green light emitted from the screens. Visual stimulus protocol was as reported previously (Kubo et al., 2014). In each experiment session, gratings moved horizontally in eight phases (3 s each at spatial frequency of 0.033 cycles/degree and temporal frequency of 2 cycles/sec, interspersed with 10 s stationary gratings). Four of the eight phases are monocular, four are binocular: 1) left nasalward, 2) left temporalward, 3) right temporalward, 4) right nasalward, 5) clockwise, 6) counter-clockwise, 7) forward, 8) backward. The sequence of eight phases was repeated three times. During visual stimulation, GCaMP fluorescence was imaged at about 3 Hz (0.501 µm/pixel, laser power ca. 15 mW after the objective, imaging region of about 256 x 256 µm). We typically imaged a volume centered around the pretectal neuropil with the z-step size of 10 μm.

#### Calcium imaging analysis

To eliminate motion artifacts, raw GCaMP image time series was registered to median image of all frames using TurboReg ImageJ plugin. To identify neurons that show correlated activity to visual stimuli, we performed regressor-based correlation analysis as described previously (Kubo et al., 2014; Miri et al., 2011). Briefly, the visual stimulus data and the registered image time series were loaded using a custom MATLAB script. To detect cells that respond to optic flow stimuli, we generated a motion-sensitive variable that is 1 during motion phases and 0 during motionless phases of the visual stimuli. This variable was then convolved with a kernel of GGCaMP6s kinetics (τdecay = 3 s for GCaMP6s and τdecay = 4 s for H2B-GCaMP6s) to generate a motion-sensitive variable, called regressor. Based on the correlation value with the regressor in each pixel of the time series, a heat map (Z-score map) was generated and we selected individual cells (ROIs) for a subsequent correlation analysis. Care was taken to only include individual cell-like structures and avoid neuropil signals.

Response type classification was performed as previously described (Kubo et al., 2014). In short, since there were 8 different stimulus phases in the visual stimulus, a total of 2^8 (256) binary response patterns were conceivable. In order to classify the response of each cell as a particular response type, 256 convolved regressors were generated that corresponded to the 2^8 possible combinations of monocular and binocular stimulations in nasalward and temporalward direction. The regressor that showed the highest correlation with the cell’s response (best regressor) defined the response type of the cell. Average response was calculated by averaging ΔF/F during each of the 8 motion phases across the 3 repetitions.

#### Cell ablations

Larvae expressing the enhanced potency nitroreductase (epNTR)-mScarlet were sorted at 3 dpf. Cell ablation was induced at 5 dpf by incubating larvae with 2 mM metronidazole (Mtz) for 24 hours, followed by washout and recovery with E3 medium for more than 20 hours. Behavioral experiments were carried out at 7 dpf. Successful ablation was confirmed by confocal imaging before and after treatment with Mtz.

#### Behavior assays for Optokinetic response (OKR), optomoter response (OMR) and spontaneous swimming

We adapted previously established methods for OKR assays (Schoonheim et al., 2010) with some modifications. Sinusoidal grating stimuli were generated by a custom-written script using PsychoPy and presented to a custom-built oLED arena that consists of four oLED screens (3.81 inch, Wisecoco) that covered 360 degrees around the fish. Binocular gratings moved horizontally in clockwise and counter-clockwise directions (duration: 60 s at spatial frequency of 0.033 cycles/degree and temporal frequency of 2 cycles/sec, interspersed with 60 s stationary gratings), with 2 times repetitions. Eye movements were recorded at 10 frames per second (fps) using a CMOS camera (Mako U-051B, Allied Vision). Angles of left and right eyes were calculated using a custom ImageJ macro based on the “Moment Calculator” plug-in. The “OKR index” was calculated by counting the saccades in the expected direction, with CCW saccades during the CW motion for the right eye and CW saccades during the CCW motion for the left eye, focusing on those with an amplitude greater than 15 degrees. OMR was measured using previously established methods (Orger et al., 2004) with some modifications. A group of 6 larvae were placed in a custom-built acrylic chamber (2 x 2 x 25 cm, RIKEN machine shop). The chambers were placed on a 23-inch monitor, on which visual stimulus was presented. Fish behavior was recorded at 10 fps using a CMOS camera (CS505MU, Thorlabs). Before the OMR tests, animals were kept in the chamber for at least 2.5 minutes to allow habituation to light and temperature conditions of the experiment. Sinusoidal grating stimuli were generated by a custom-written script using PsychoPy. Moving gratings (grating width of 1 cm and temporal frequency of 1 cycle/sec) were in motion for 30 s. Average distance traveled was calculated by the difference between the average positions of 6 fish in one chamber before and after the moving grating stimulus. To measure spontaneous swimming, one larva was placed in a plastic weighing dish (70 x 70 x 22 mm, BIO-BIK). After habituation for 3 min, swimming was recorded for 1.5 min at 10 fps. Swimming trajectories were tracked using ImageJ Plugin TrackMate (Ershov et al., 2022).

### QUANTIFICATION AND STATISTICAL ANALYSIS

To assess the normality of behavioral data, a Q-Q plot and the Shapiro-Wilk normality test were performed. If normality was confirmed, Welch’s two-sample t-test was used (OMR, spontaneous locomotor activity). For data that did not follow a normal distribution, the Mann-Whitney U test was applied (OKR). All statistical analyses were conducted using R (version 4.1.0).

### DATA AND SOFTWARE AVAILABILITY

Data and software will be made available upon request.

## Notes

### Competing Interest Statement

The authors have declared no competing interest.

