## Supplemental Data for "Molecular and functional dissection using CaMPARI-seq reveals the neuronal organization for dissociating optic flow-dependent behaviors"

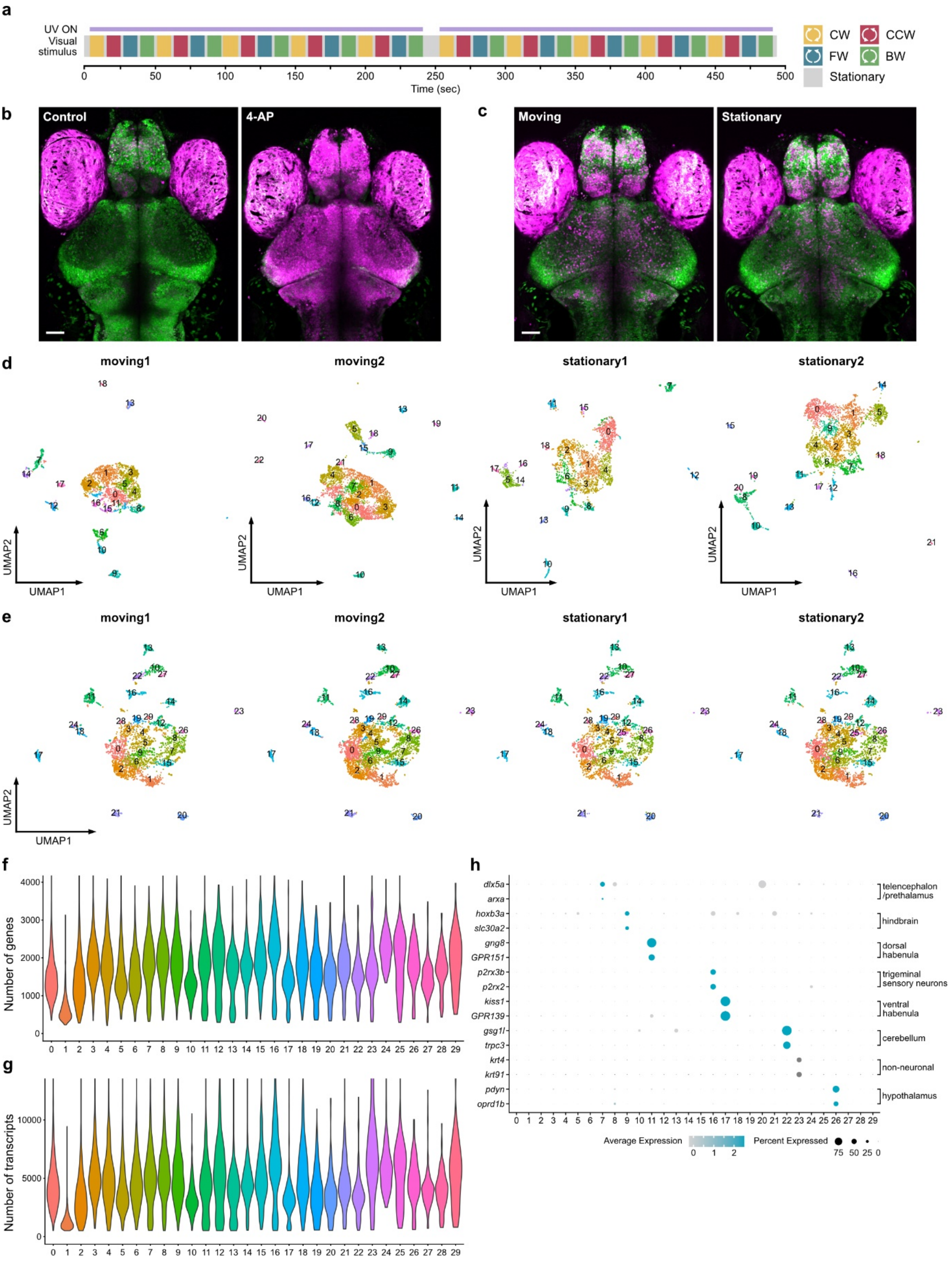


**Extended Data Fig. 1 Strategies for identifying marker genes for optic flow-responsive pretectal neurons.**

**a,** Protocol for CaMPARI2 photoconversion during optic flow stimulation. CW, clockwise; CCW, counter-clockwise; FW, forward; BW, backward. **b,** Validation of NLS-CaMPARI2 photoconversion in *Tg(elavl3:NLS-CaMPARI2)* larvae. Larvae were illuminated with UV at 405 nm either in the absence or presence of the 4-aminopyridine (4-AP). **c,** Photoconversion of CaMPARI2 during optic flow stimulation using Stationary or Moving gratings. Photoconversion and visual stimulation were performed as shown in (**a**). **d,** UMAP embedding of two replicates from Moving condition (moving1, moving2) and two replicates from Stationary conditions (stationary1, statioary2) before the integration. Clustering was performed within each dataset. **e,** As in (**d**), but after the integration. Clustering was performed on the integrated dataset. **f,g,** Violin plots showing the distribution of the number of transcripts detected per cell (**f**) and the number of genes detected per cell (**g**) in each cluster. **h,** Marker genes and annotation for Clusters 7, 9, 11, 16, 17, 22, 23 and 26, which were excluded from the further analysis. Scale bars, 50 μm.

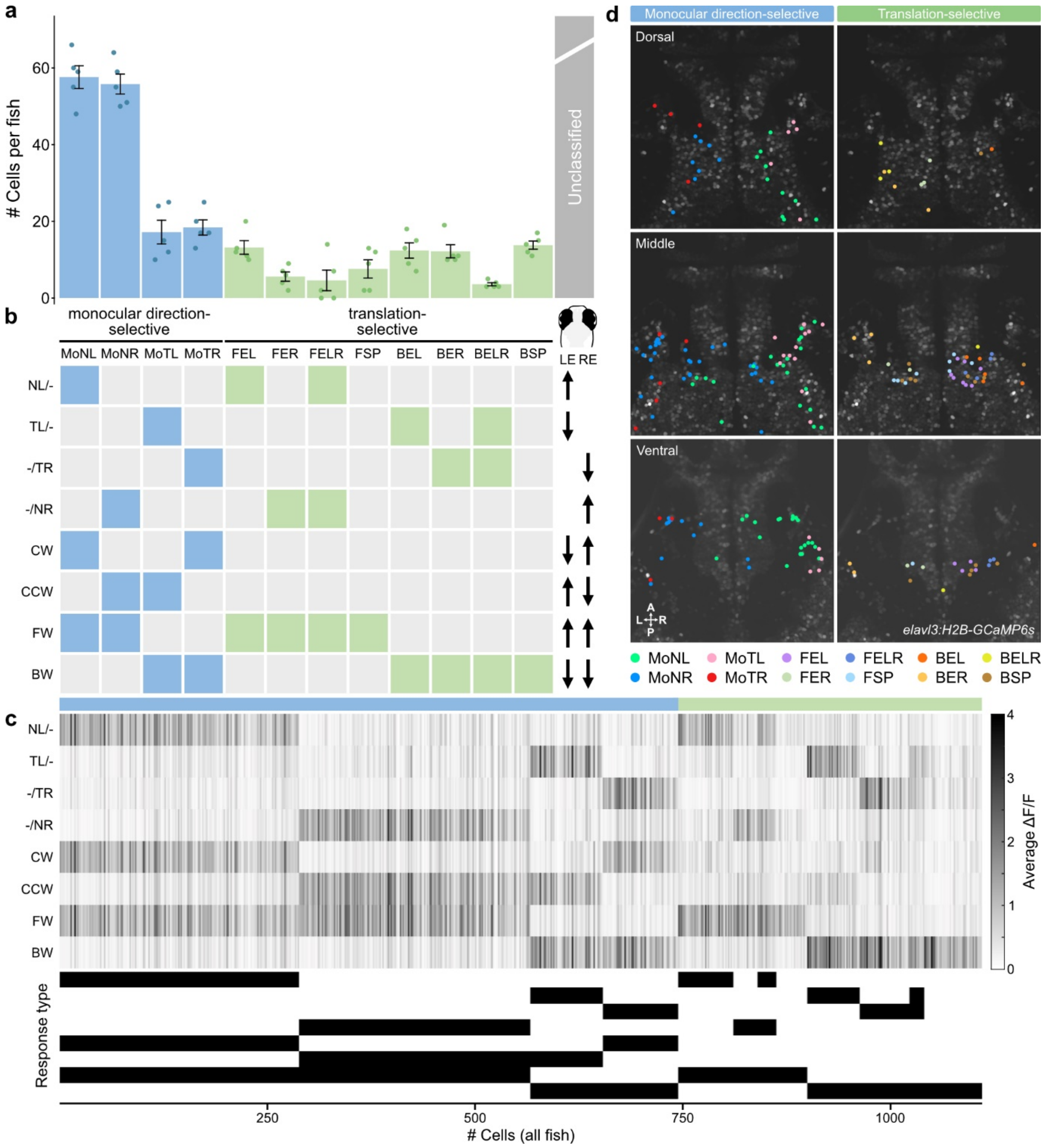


**Extended Data Fig. 2. Frequency of monocular and binocular optic flow-responsive cells recorded in the pan-neuronal *Tg(elavl3:H2B-GCaMP6s)* line.**

**a,** A histogram of the frequent response types (“simple”, blue, “translation-selective”, green) detected in *Tg(elavl3:H2B-GCaMP6s)* larvae ($n$ = 5 fish). The remaining response types were listed as unclassified (gray). Each bar represents the average # of cells per fish. Error bars represent ±SEM. **b,** Nomenclature of the visual stimulus protocol and response types as previously described (Kubo et al., 2014). **c,** (Top) Raster plot showing the responses to the eight stimulus phases ($n$ = 1110 cells, pooled from 5 fish). Cells are ordered according to their correlation coefficient to the corresponding regressor (within each response type). ΔF/F values across the 3 repetitions of the visual stimulation were averaged. (Bottom) Binary response type barcode of each cell. **d,** Spatial distribution of the response types (as shown in (**a**)) in a representative *Tg(elavl3:H2B-GCaMP6s)* fish. Cells are color-coded according to their response type. Average image of GCaMP6s signal of the dorsal, middle and ventral planes of the pretectum is used as a background. For the middle and ventral planes, cells detected in the same plane, as well as those detected in 10 μm above and 10 μm below the corresponding plane, are plotted. For the dorsal plane, cells detected in the same plane, as well as those detected in 10 μm above the corresponding plane, are plotted. For the data from all individual planes, see also Extended Data Fig. 4 and 5.

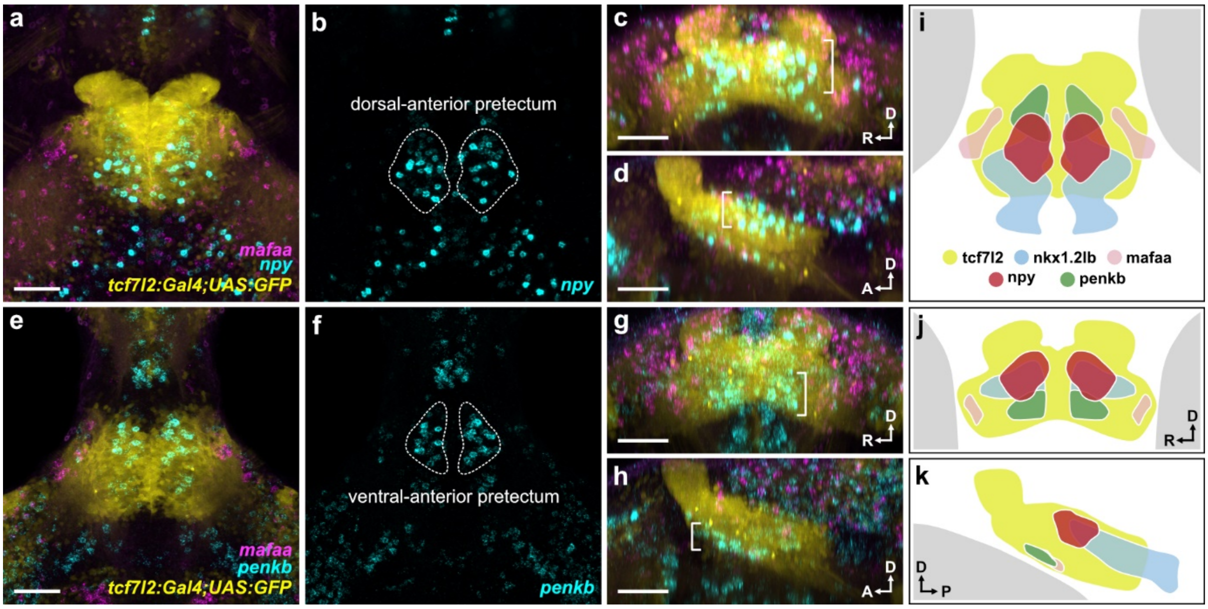


**Extended Data Fig. 3. Expression pattern of *npy*, *penkb* in the pretectum.**

**a-d,** Substack maximum z-projections of double HCR-FISH stains of *npy* and *mafaa* in a *Tg(tcf7l2-hs:Gal4FF);Tg(UAS:GFP)* larva in the dorsal (**a**,**b**), coronal (**c**) and sagittal (**d**) views. **e-h,** Substack maximum z-projections of double HCR-FISH stains of *penkb* and *mafaa* in *Tg(tcf7l2-hs:Gal4FF);Tg(UAS:GFP)* larva in the dorsal (**e**,**f**), coronal (**g**) and sagittal (**h**) views. **i-k,** Schematic of the expression patterns in the pretectum in the dorsal (**i**), coronal (**j**) and sagittal (**k**) views. D, dorsal; A, anterior; R, right; P, posterior. Scale bars, 50 µm.

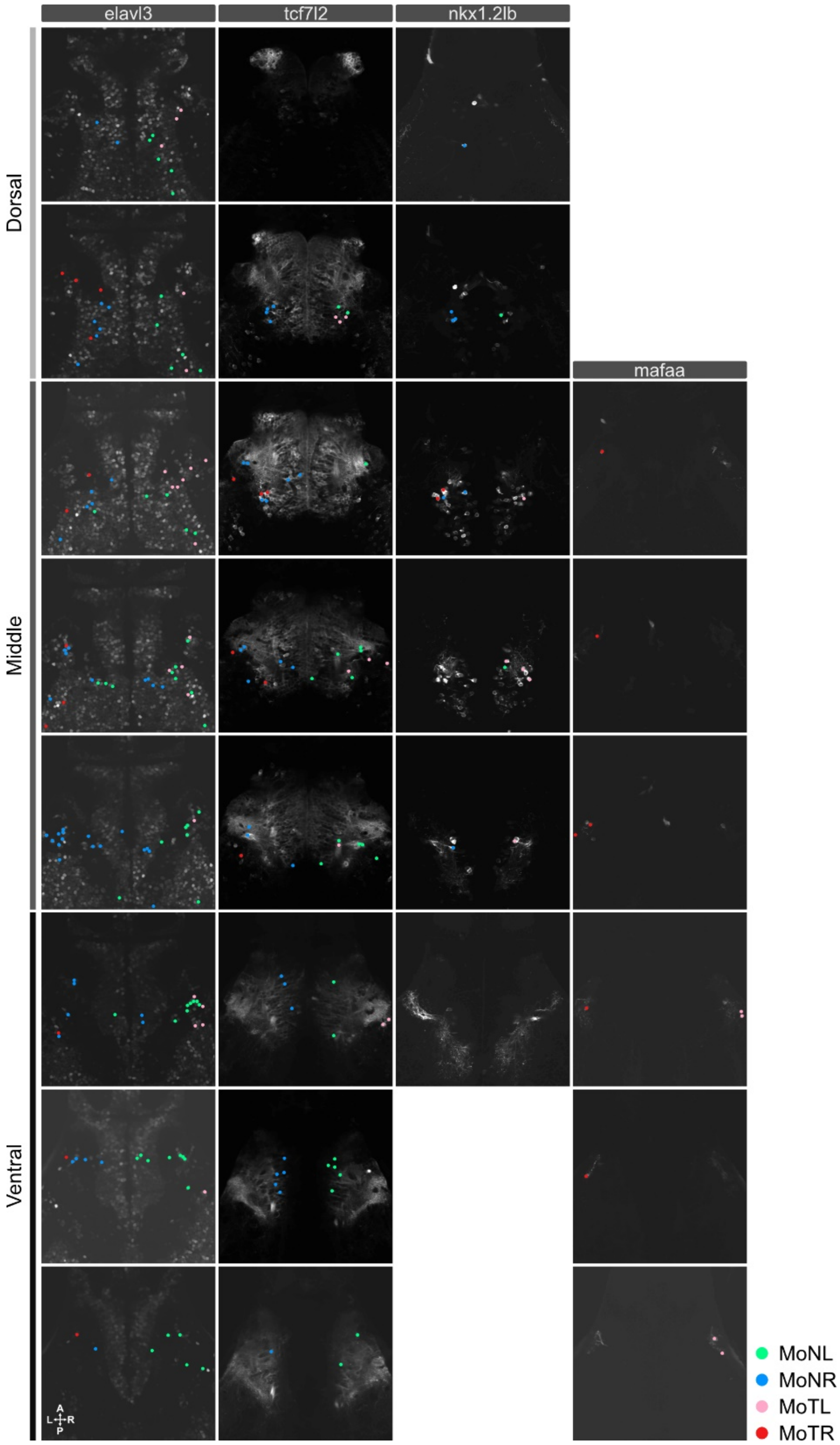


**Extended Data Fig. 4 Spatial distribution of monocular response types in *elavl3*-, *tcf7l2*-, *mafaa*- and *nkx1.2lb*-positive neurons.**

Spatial distribution of 4 monocular response types detected in *Tg(elavl3:H2B-GCaMP6s)*, *Tg(mafaa-hs:Gal4FF);Tg(UAS:GCaMP6s), Tg(nkx1.2lb-hs:Gal4FF);Tg(UAS:GCaMP6s)* and *Tg(mafaa-hs:Gal4FF);Tg(UAS:GCaMP6s)* fish. A representative larva from each transgenic line is shown.

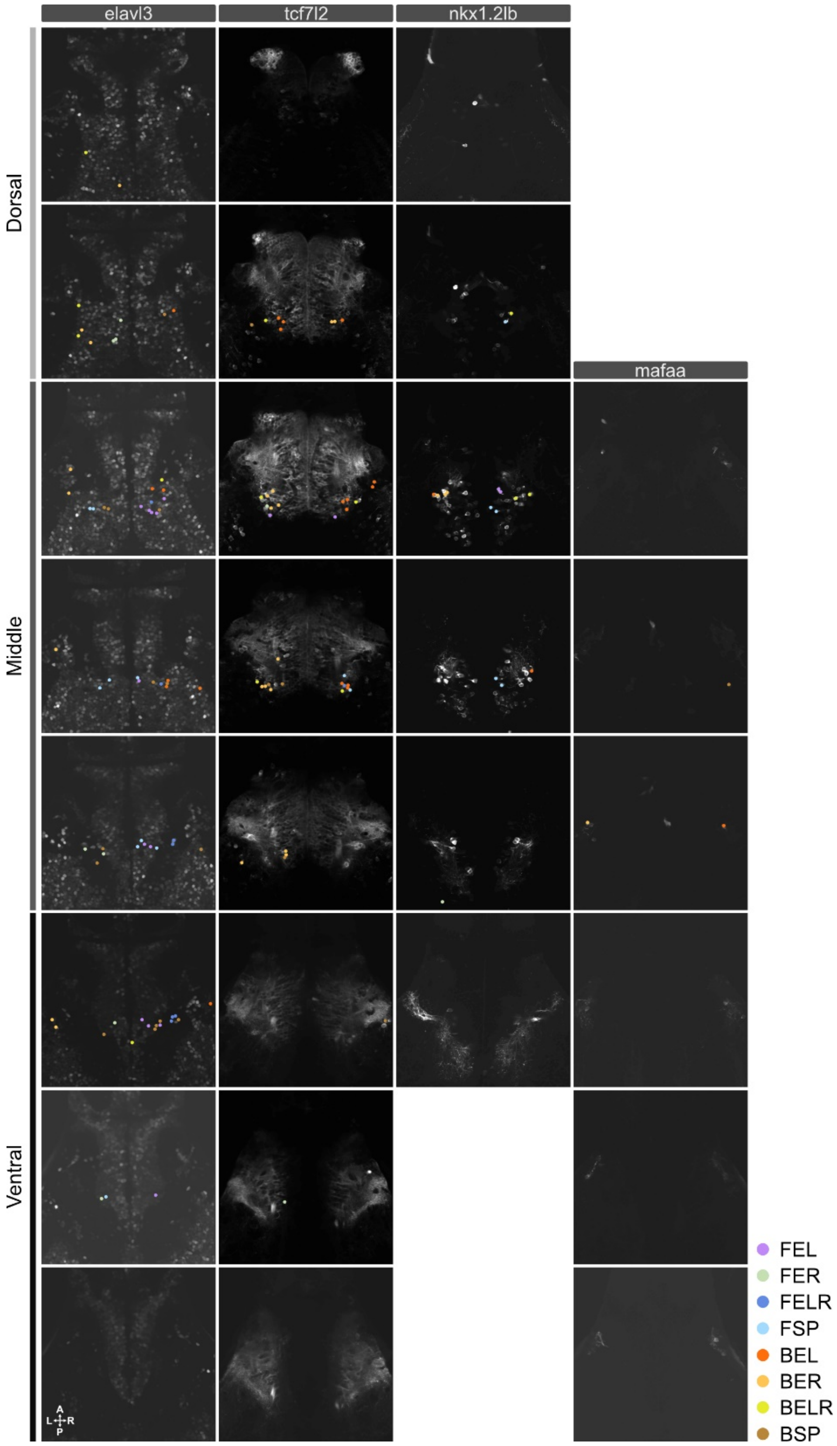


**Extended Data Fig. 5 Spatial distribution of translation-selective response types in *elavl3*-, *tcf7l2*-, *mafaa*- and *nkx1.2lb*-positive neurons.**

Spatial distribution of translation-selective response types detected in *Tg(elavl3:H2B-GCaMP6s)*, *Tg(mafaa-hs:Gal4FF);Tg(UAS:GCaMP6s)*, *Tg(nkx1.2lb-hs:Gal4FF);Tg(UAS:GCaMP6s)* and *Tg(mafaa-hs:Gal4FF);Tg(UAS:GCaMP6s)* fish. A representative larva from each transgenic line is shown.

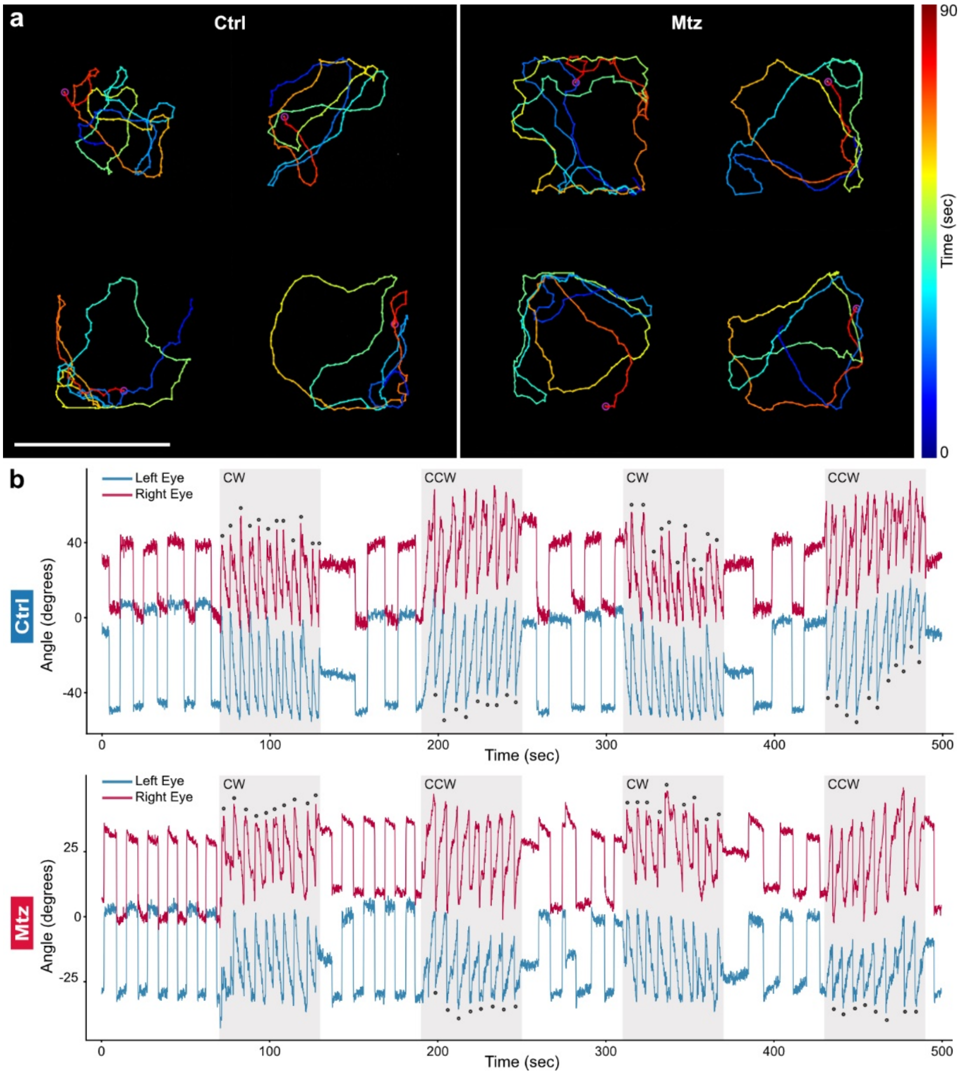


**Extended Data Fig. 6 Behavioral phenotype of *nkx1.2lb*+ pretectal neurons ablated fish.**

**a,** Locomotor activity of control (Ctrl) and ablated (Mtz) fish using *Tg(nkx1.2lb-hs:Gal4FF);Tg(UAS-hs:loxp-TagBFP-loxp-epNTR-mScarlet);Tg(tcf7l2-hs:Cre)* larvae. Trajectories from 4 example larvae during 90 second are shown for control and Mtz-treated larvae. **b,** Example eye traces of control and Mtz-treated larvae. Binocular clockwise (CW) and counter-clockwise (CCW) grating stimuli are indicated as gray shadings. Saccades are indicated by dots.
